# Bipartite anchoring of SCREAM enforces stomatal initiation by coupling MAP Kinases to SPEECHLESS

**DOI:** 10.1101/587154

**Authors:** Aarthi Putarjunan, Jim Ruble, Ashutosh Srivastava, Chunzhao Zhao, Amanda L. Rychel, Alex K. Hofstetter, Xiaobo Tang, Jian-Kang Zhu, Florence Tama, Ning Zheng, Keiko U. Torii

## Abstract

Cell-fate in eukaryotes is regulated by MAP Kinases (MAPKs) that translate external cues to cellular responses. In plants, two MAPKs, MPK3/6, regulate diverse processes of development, environmental response, and immunity. Yet, the mechanism bridging these shared signaling components with a specific target remains unresolved. Focusing on the development of stomata, epidermal valves for gas exchange and transpiration, we report here that the bHLH protein SCREAM functions as a scaffold by recruiting MPK3/6 to downregulate SPEECHLESS, a transcription factor initiating stomatal cell lineages. SCREAM directly binds with MPK3/6 through an evolutionarily-conserved yet unconventional bipartite motif. Mutations in this motif abrogate association, phosphorylation and degradation of SCREAM, unmask hidden non-redundancies between MPK3 and MPK6, and result in uncontrolled stomatal differentiation. Structural analyses of MPK6 at the 2.75Å resolution unraveled bipartite binding of SCREAM with MPK6, that is distinct from an upstream MAPKK. Our findings elucidate, at the atomic resolution, the mechanism directly linking extrinsic signals to transcriptional reprogramming during the establishment of stomatal cell-fate, and highlight a unique substrate-binding mode adopted by plant MAPKs.

## Introduction

Organized differentiation of functional tissue types is a critical step towards ensuring survival and the overall fitness of multicellular organisms. Fundamental to these processes are mitogen-activated protein kinases (MAPK) cascades, consisting MAP3K (MAPKKK or MEKK), MAP2K (MAPKK or MEK), and MAPK that phosphorylates and activates downstream targets to regulate cell proliferation, differentiation, and polarity ^1^. In plants, the MAPK cascade is known to broadly influence development, environmental response, and immunity ^2-5^. Among the 20 known MAPKs in *Arabidopsis*, MPK3 and MPK6 play a predominant role in diverse developmental programs, including embryo patterning, inflorescence architecture, floral abscission, anther and ovule development, and stomatal patterning ^6-10^. It is thus imperative to understand how these MAPKs recognize their target substrates and specifically activate individual developmental programs. Despite the recently reported partial crystal structure of MPK6 ^11^, the structural bases for its substrate association remains unclear.

Development of stomata, cellular valves on the plant epidermis essential for efficient gas exchange and water control, is an excellent system to understand how external signals are interpreted for the specification of cell fate ^12, 13^. The EPIDERMAL PATTERNING FACTOR (EPF) family of upstream peptide signals are secreted from stomatal precursors and perceived by the ERECTA-family receptor-like kinases in their neighboring cells. This activates the downstream MAPK cascade composed of YODA (YDA) MAPKKK, two redundant MAPKKs - MKK4/5, and two redundant MAPKs - MPK3/6, which culminates with the prevention of stomatal differentiation by phosphorylation and inhibition of the bHLH transcription factor, SPEECHLESS (SPCH) ^10, 14-17^. Recent studies have shown that BREAKING OF ASYMMETRY IN THE STOMATAL LINEAGE (BASL) recruits YDA and MPK3/6 to the cortical polarity site ^18^. This leads to reduced SPCH accumulation and eventual loss of stomatal fate in one of the two daughter cells of stomatal precursors ^19^. However, unlike the *mpk3 mpk6* double mutant, which produces an epidermis solely composed of stomata ^10^, the *basl* null mutant produces a nearly normal epidermis, with occasional paired stomata due to mis-specification of asymmetric cell divisions ^20^. Thus, the action of BASL cannot explain the mechanism by which the MAPK cascade enforces the decision to initiate stomatal differentiation.

In search of an elusive factor that recruits MAPKs to the nucleus to down-regulate SPCH, we revisited the gain-of-function mutant of *SCREAM* (*SCRM;* also known as *ICE1*), *scrm*-D, which confers constitutive stomatal differentiation (Kanaoka et al., 2008). SCRM and its paralog SCRM2 function as partner bHLH proteins for SPCH as well as for the later-acting stomatal bHLH proteins, MUTE and FAMA ^21-23^. The scrm-D protein has an R-to-H amino-acid substitution within its conserved motif, named the KRAAM motif ^21^. Yet, nothing is known about its exact function.

Here we report that the bHLH protein SCRM physically bridges MAPKs and SPCH and plays a direct role in enforcing entry into the stomatal lineage. A bipartite module with the unique KRAAM motif, in conjunction with an upstream conventional MAPK docking motif, mediate specific interactions with MPK3 and MPK6 in the stomata development pathway. Precise dissection of the binding interface revealed distinct SCRM-binding properties of MPK3 and MPK6, and further unmasked cryptic functional non-redundancies between these two redundant MAPKs. Structural analysis of MPK6 at 2.75Å resolution, together with *ab initio* modeling of the MPK6-SCRM protein-peptide complex, identified the exact amino-acid residues serving the binding interface. Our work provides the mechanistic basis for SCRM to function as an integrator of upstream repressive cues and downstream activators during stomatal development, and highlights the unique recruitment mechanism of the plant MAPK signaling cascade.

## Results

### SCRM serves as a scaffold to recruit MAPK to SPCH

MPK3/6-mediated phosphorylation of SPCH determines the entry into the stomatal cell lineage (Fig. 1a) ^10^. It remains unknown, however, whether SPCH directly associates with MPK3/6. To address this question, we first performed yeast two-hybrid (Y2H) assays with SPCH^ΔN^ (truncated SPCH without its N-terminal extension to overcome auto-activation) in pairwise combinations with SCRM, MPK3, and MPK6 (Fig. 1b). No interactions of SPCH^ΔN^ with MPK3 or MPK6 were detected, whereas SPCH^ΔN^ heterodimerizes with SCRM (Fig. 1b). This result suggests that the interaction between SPCH and MPK3/6 is either too weak or too transient to be detected, requires the SPCH N-terminal domain, or that it otherwise requires SCRM as a scaffolding partner.

**Fig. 1:**
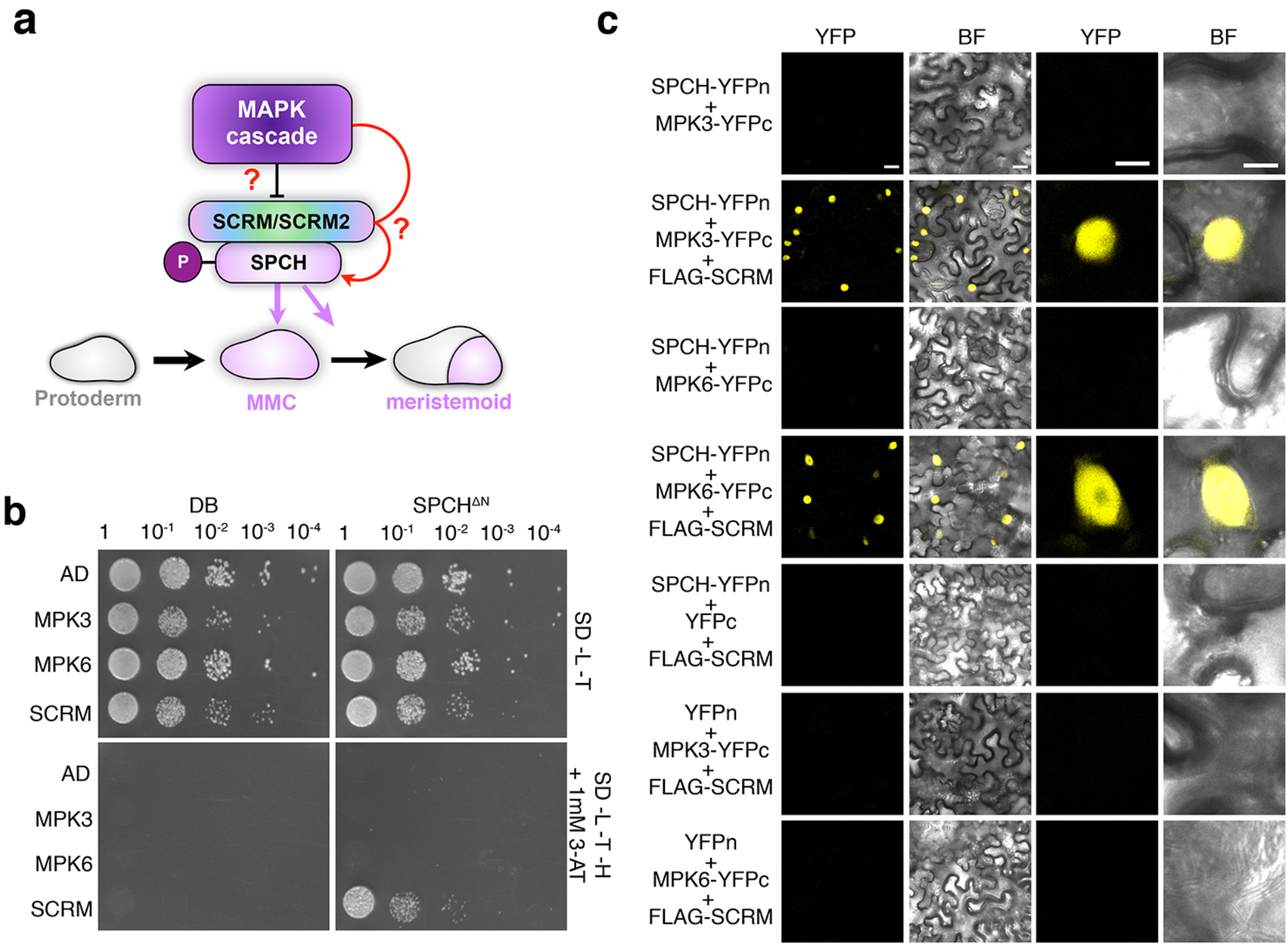
SCRM serves as a scaffold to recruit MAPK to interact with and modulate the stability SPCH. **a,** Schematic illustrating the influence of the MAPK cascade in regulating cell fate specification with the direct connection yet to be demonstrated (question mark). MMC, meristemoid mother cell. **b,** Yeast two-hybrid (Y2H) assays. Bait: the DNA binding domain alone (DB) and SPCH(ΔN-term). Prey: the activation domain alone (AD), MPK3, MPK6, SCRM and scrm-D. No detectable interaction between MPK3/MPK6 and SPCH is detected. **c,** Bimolecular Fluorescent Complementation (BiFC) assays. Shown are 3-week old *N. benthamiana* leaves agroinfiltrated using pairwise combinations of SPCH-YFPn and MPK3-YFPc, MPK6-YFPc along with 35S:: FLAG-SCRM (Scale bars = 25 μm). Right two panels are magnified images of a representative nucleus (scale bars = 10 μm). SPCH does not interact with MPK3/6 by itself but interacts with them only in the presence of SCRM.

To test these hypotheses, we designed a ‘three-way’ bimolecular fluorescent complementation (BiFC) assays. First, the full-length SPCH protein was fused to the N-terminal half of YFP (YFPn) and MPK3/6 were fused to the complementary half YFP (YFPc). When co-expressed in *N. benthamiana* leaves, no signal was observed, indicating that SPCH does not directly interact with MPK3/6 (Fig. 1c). A subsequent immunoblot analysis detected the accumulation of SPCH protein (Fig. S1), thus refuting the possibility that the lack of YFP signal is due to the absence of SPCH protein accumulation. SPCH forms a heterodimer with SCRM to initiate stomatal differentiation (Fig. 1a) ^21, 23^. We next introduced non-fluorescently tagged SCRM (FLAG-SCRM driven by its own promoter) along with SPCH-YFPn and MPK3/6-YFPc. Only in the presence of SCRM, SPCH and MAPKs are able to reconstitute strong YFP signals in the nuclei of *N. benthamiana* (Fig. 1c). We thus conclude that SCRM serves as the scaffold to couple MPK3 and MPK6 with SPCH.

### The evolutionarily conserved SCRM KiDoK motif defines a direct MPK3/6 interaction surface

To elucidate how scaffolding of SCRM with MPK3/6 and SPCH translate into organized differentiation of stomatal-lineage cells, we revisited the *SCRM* gain-of-function mutant, *scrm*-D ^21^. The ‘stomata-only’ epidermis in *scrm-D* is essentially identical to that observed by the loss of *MPK3* and *MPK6* (*mpk3 mpk6*) ^10^. Sequence analyses of SCRM and its orthologs revealed the presence of a putative MAP Kinase Docking site immediately upstream of the KRAAM motif, which is mutated to KHAAM in scrm-D (Figs. 2a, S2a). To understand the role of the conserved region encompassing the MAP <u>Ki</u>nase <u>Do</u>cking and <u>K</u>RAAM (KiDoK) motif, we first used it as a bait and performed an unbiased Y2H screen against an *Arabidopsis* seedling cDNA library (see Methods). From 158 million interactions tested, only two proteins, MPK3 and MPK6, were identified with confidence scores high enough to be designated as real interactors (Table S1). Further targeted Y2H assays validated the interaction between the SCRM KiDoK motif and MPK3 and MPK6, but not with the distantly-related AtMAPK homolog, MPK4, that has no major role in stomatal development (Figs. 2b, S2b). Strikingly, the *scrm-D* mutation in the KiDoK motif completely abolished interaction with both MPK3 and MPK6 (Fig. 2b). We subsequently performed BiFC assays in *N. benthamiana* ^24^ (Fig. 2c). Strong YFP signals were reconstituted in the nuclei of leaves co-expressing SCRM-YFPn with MPK3 or MPK6 fused with a complementary YFPc. In contrast, no signal was detected in the pairwise combination of scrm-D-YFPn with MPK3-YFPc or MPK6-YFPc. These results indicate that the scrm-D (R236H) substitution abolishes the direct interaction with MPK3/6.

**Fig. 2:**
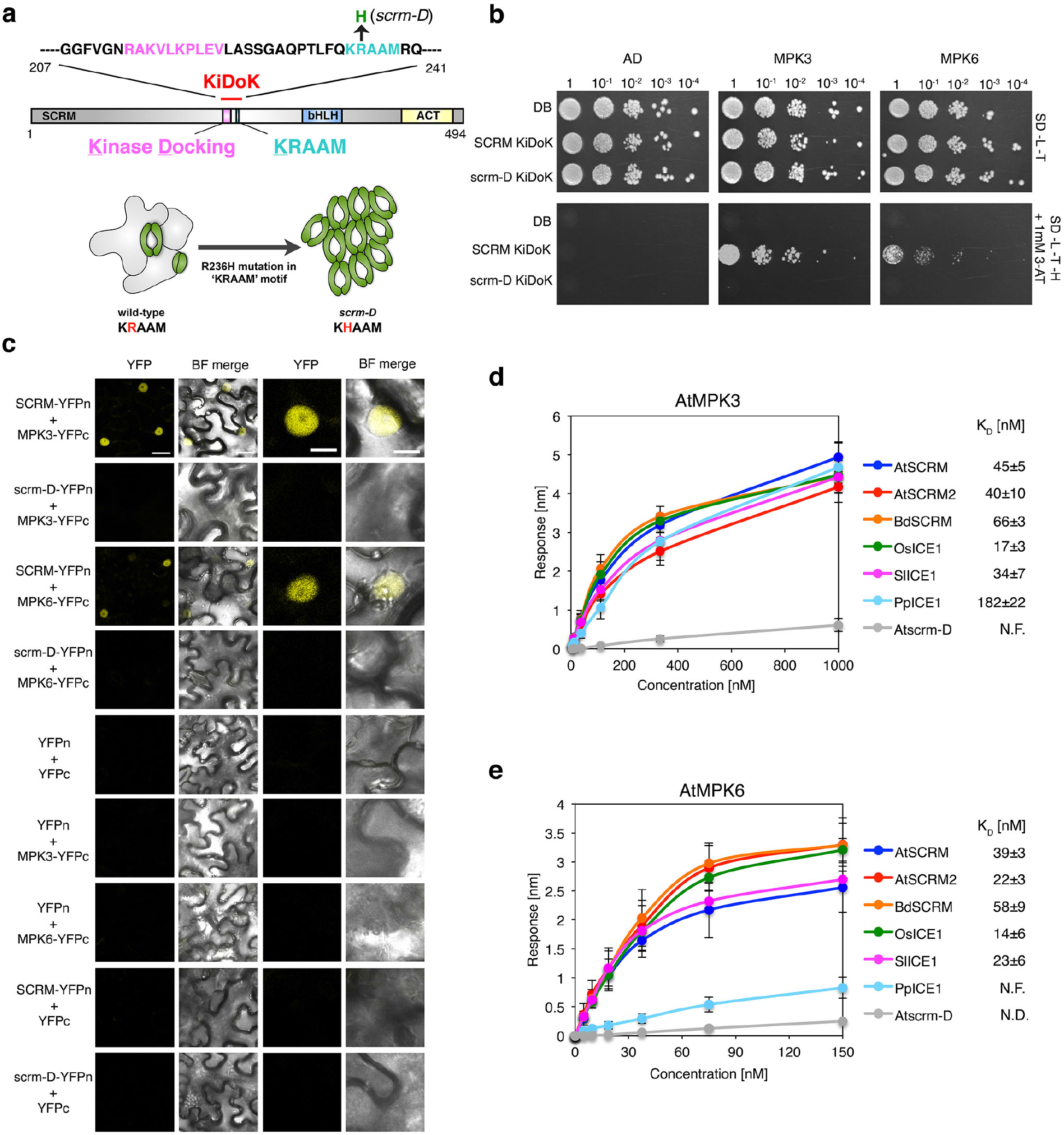
The evolutionarily-conserved KiDoK motif of SCRM defines a direct MPK3/6 interaction surface. **a,** Diagram of the SCRM/ICE1 protein. Pink, predicted Kinase Docking motif; cyan, the KRAAM motif; blue, the bHLH domain; yellow, the C-terminal domain. Shown above is the amino-acid sequence of the Kinase Docking motif in pink, KRAAM motif in cyan, and the scrm-D mutation (R236H) in green. Below, is a cartoon representation of the wild-type and the *scrm-D* epidermis. **b,** Yeast two-hybrid (Y2H) assays. Bait: DNA-binding domain alone (DB), the wild-type KiDoK motif (SCRM KiDoK) and its scrm-D version (scrm-D KiDoK); Prey: activation domain alone (AD), MPK3, and MPK6. **c,** BiFC assays. Shown are 3-week old *N. benthamiana* leaves agroinfiltrated with pairwise combinations of SCRM-YFPn and scrm-D-YFPn with MPK3-YFPc and MPK6-YFPc. Scale bars = 25 μm. Right two panels are magnified images of a representative nucleus. Scale bars = 10 μm. **d, e,** Quantitative analyses of SCRM KiDoK from various plant species with MPK3 (**d**) as well as MPK6 (**e**) using bio-layer interferometry (BLI). Shown are *in vitro* binding response curves for GST-MPK3 and biotinylated peptides of the KiDoK motif from AtSCRM, BdSCRM, OsSCRM, AtSCRM2, PpSCRM, Atscrm-D and SlSCRM, respectively, at six different concentrations (1000, 333.33, 111.11, 37.04, 12.34, 4.11 nM). The response curves for GST-MPK6 with the same biotynylated peptides were obtained by performing the experiment at lower concentrations (150, 75, 37.5, 18.75, 9.375, 4.6875 nM) to enable a better fit for calculating K_D_ values. Values represent the mean and error bars the SD of three independent experiments performed. K_D_ values are listed next to the graph legend.

The KiDoK motif of SCRM is highly conserved amongst vascular and non-vascular land plants (Fig. S2) ^25^. To explore whether the SCRM KiDoK motif constitutes an interaction module for MPK3/6 in diverse land plant lineages, we performed *in vitro* direct binding and quantitative kinetic assays of the KiDoK motif peptide from dicots (Arabidopsis *SCRM* and *SCRM2*, and tomato *SIICE1*), monocots (rice *OsICE1* and *Brachypodium BdSCRM*) and a non-vascular plant (*Physicomitrella PpICE*) with purified AtMPK3 and AtMPK6 proteins using BioLayer Interferometry (BLI) (Figs. 2d, e). *BdSCRM2* lacks the KiDoK motif altogether, and thus was not used for the analyses (see Discussion). All orthologous KiDoK motif peptides showed tight binding with MPK3, with K_D_ values at nanomolar levels, whereas all but PpICE1 exhibited tight binding to MPK6 (Figs. 2d, e). Again, scrm-D did not show any appreciable affinity to MPK3 or MPK6. Our results establish the SCRM KiDoK motif as an evolutionarily conserved direct interaction surface for MPK3/6.

### Functional analysis of the SCRM KiDoK motif unmasks the basis of non-redundancies of MPK3 and MPK6

To address the functional significance of the SCRM KiDoK motif, we performed site-directed mutagenesis of the K235 and/or R236 residues in the KRAAM motif to a non-charged residue, alanine (KRAAM→KAAAM, AAAAM, ARAAM, AHAAM), as well as deleted the Docking motif from both SCRM and scrm-D. We then performed interaction analyses of these mutant variants with MPK3/6 using Y2H assays. Whereas the various alanine substitutions in the KRAAM motif abolished interaction with both MPK3 and MPK6, SCRM_ΔDocking motif only abolished interaction with MPK6, but not with MPK3 (Fig. 3a).

**Fig. 3:**
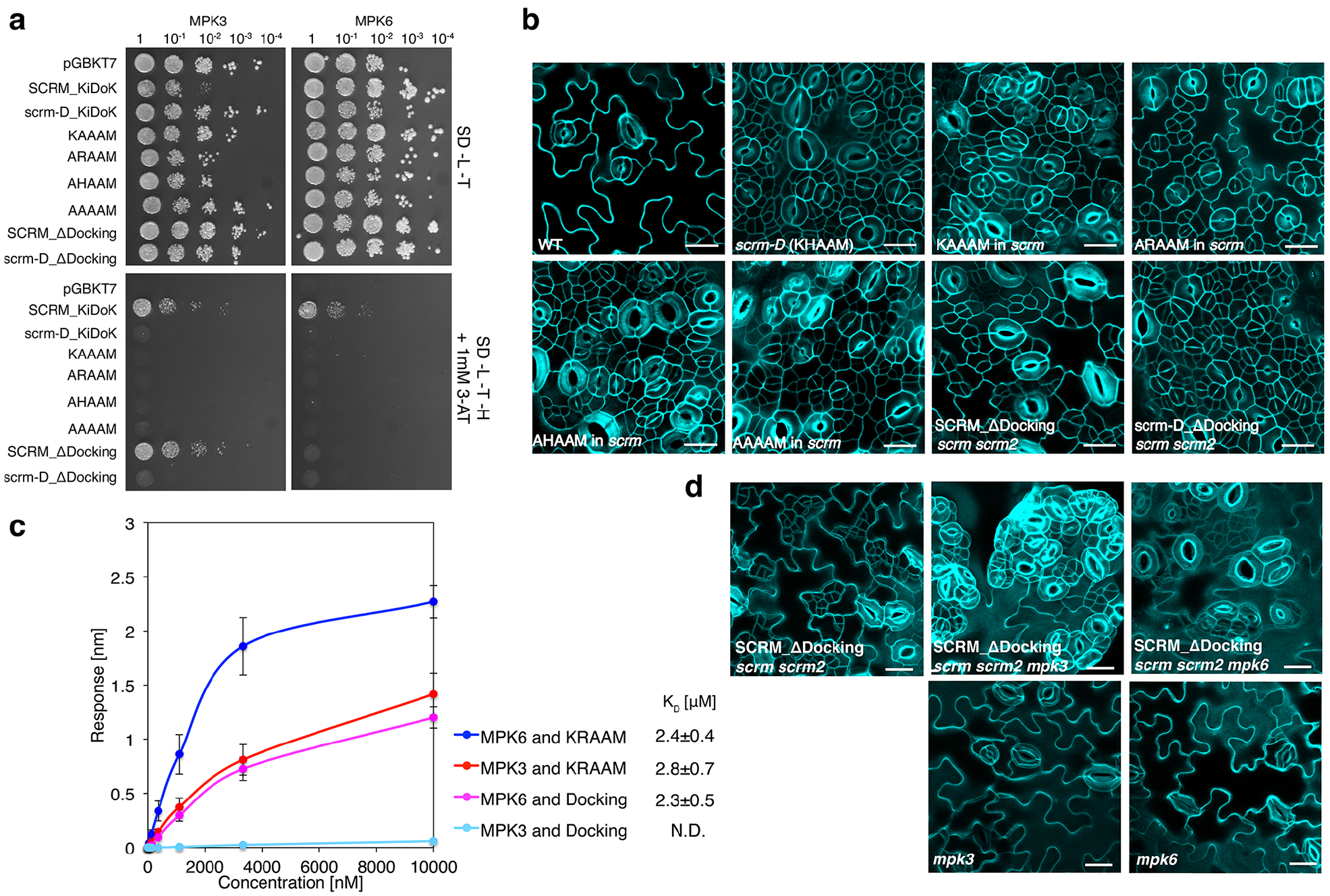
MPK3 and MPK6 exhibit different binding modes to the SCRM KiDoK motif to repress stomatal cell fate. **a,** Y2H assays of SCRM-KiDoK motif substitutions/deletions with MPK3/6. Bait: the DNA-binding domain alone (DB), the wild-type KiDoK motif (SCRM_KiDoK), scrm-D version of the KiDoK motif (scrm-D_KiDoK), ‘KAAAM’, ‘ARAAM’, ‘AHAAM’, and ‘ARAAM’ versions of the SCRM KiDoK motif, and Kinase-Docking deletion in SCRM (SCRM_ΔDocking) and scrm-D (scrm-D_ΔDocking). Prey: the activation domain fused to MPK3 and MPK6. SCRM_ΔKiDoK motif only abolished interaction with MPK6, but not with MPK3. **b,** Cotyledon abaxial epidermis of 7 day-old wild-type (WT) and transgenics Arabidopsis seedlings expressing the SCRM-KiDoK motif substitutions/deletions (*scrm-D, SCRMpro::SCRM-‘KAAAM’, SCRMpro::SCRM-‘ARAAM’, SCRMpro::SCRM-‘AHAAM’,* and *SCRMpro::SCRM-‘AAAAM’* in an *scrm* background, and *SCRMpro::FLAG-SCRMΔDocking* and *SCRMpro::FLAG-scrm-DΔDocking* in *scrm scrm2*. Scale bars = 20 μm. **c,** Quantitative analysis of interactions between the KRAAM motif and the Kinase Docking motif of SCRM with GST-MPK3 and GST-MPK6 using BLI. Shown is the *in-vitro* binding response curve for purified GST-MPK3/6 and biotinylated KRAAM motif peptide and Docking motif peptide at 6 different concentrations (10000, 3333.33, 1111.11, 370.4, 123.4, 41.1 nM). Values represent the mean and error bars the S.D. of three independent experiments performed. K_D_ values are listed next to the graph legend. **d,** Representative stomatal phenotypes of *SCRMpro::3x-FLAG-SCRM_ΔDocking* in *scrm scrm2, scrm scrm2 mpk3* and *scrm scrm2 mpk6* along with *mpk3* and *mpk6* single mutants. Abaxial epidermis from 7-day-old plants were imaged. Scale bars = 20 μm

To examine the *in vivo* phenotypic consequence of these mutations, we introduced each mutant construct driven by the endogenous *SCRM* promoter into the *scrm* null mutant. Like *scrm-D*, all alanine substitutions conferred severe stomatal clustering (Fig. 3b), emphasizing that loss of MPK3/6 association due to the altered KRAAM degron motif triggers constitutive stomatal differentiation. Compared to *SCRMpro::scrm-D_ΔDocking,* which confers a phenotype identical to *scrm-D* when introduced into the *scrm scrm2* double knockout background*, SCRMpro::SCRM_ΔDocking* led to a weaker phenotype (Fig. 3b), presumably reflecting the specific loss of its binding with MPK6 but not with MPK3 (Fig. 3a). To further dissect the specific association of SCRM with MPK3 and MPK6, we truncated the KiDoK peptide into one containing only the KRAAM motif and the other containing only the Docking motif. *In vitro* quantitative binding assays using BLI showed that whereas KRAAM motif binds to both MPK3 and MPK6 with high affinity (K_D_ values: 2.8±0.7 μM and 2.4±0.4 μM respectively), the Docking motif only associates with MPK6 but not with MPK3 (K_D_ values: 2.3±0.5 μM and N.D. respectively)(Fig. 3c).

It is well known that *MPK3* and *MPK6* redundantly repress stomatal development: neither the *mpk3* nor the *mpk6* single mutant confers stomatal clustering phenotype ^10^ (Fig. 3d). Our finding points to a unique mode of their association with SCRM: MPK6 requires both the Docking and the KRAAM motifs, whereas MPK3 requires only the latter (Figs. 3a, c). To unmask the differential requirements of MPK3 and MPK6 to interact with SCRM and inhibit stomatal differentiation, we further introduced *SCRMpro::SCRM_ΔDocking* into *the scrm scrm2 mpk3* and *scrm scrm2 mpk6* triple mutant backgrounds (Fig. 3d). Indeed, the additional loss of *mpk3* led to de-repression of stomatal cell-fate, giving rise to severe stomatal clustering, whereas the additional loss of *mpk6* had no discernable phenotypic enhancement (Fig. 3d). These striking phenotypic differences between *mpk3* and *mpk6* seedlings expressing *SCRM_ΔDocking* emphasize the notion that MPK3 and MPK6 feature a major difference in their binding sites (CD domain, see below) responsible for recognizing the conventional MAP kinase Docking motif within the SCRM KiDoK sequence. Thus, by detaching the Docking and KRAAM motifs, we can unmask cryptic functional non-redundancies between MPK3 and MPK6 in repressing stomatal cell fate.

### Structural analyses of SCRM_KiDoK-MAPK interaction module

The KRAAM motif is unique to SCRM/ICE1 orthologs and has otherwise not been found to be involved in binding MAPK in plants or animals. To gain structural insight into its function, we first independently determined the crystal structure of MPK6 (residues 32-395, MPK6ΔNt) at 2.75 Å resolution (Figs. S3a, S4; see Methods). MPK6 crystallized with two molecules in the asymmetric unit and displayed packing and main chain alignment closely resembling the partial structure previously determined at 3.2 Å resolution ^11^ (Figs. S3a, S4). MPK6 forms a bi-lobed structure divided between N- and C-termini, with an ATP binding pocket between the two lobes. Secondary structure components of our MPK6 structure are given the following labeling scheme: α–helices H1-H20 and β-strands S1-S10 (Figs. S3a, S4).

Mammalian MAPKs possess a ‘docking groove’ in the C-lobe that contains a highly conserved Common-Docking (CD) domain mostly composed of negatively-charged residues that bind activators (MAPKKs/MEKs), inactivators (MKPs) and substrates ^26^. In most MAPKs, these exposed, negatively charged residues (aspartic acid or glutamic acid) are involved in mediating peptide substrate binding ^27^. We found the CD domain is mostly conserved between MPK3/6 and mammalian ERK/p38 family (Fig. S3). In fact, the structure of AtMPK6 reveals the highly conserved D353 and D356 residues exposed from H18 within the CD domain (Fig. S3).

To probe the possible role of the CD domain of MPK6 in SCRM binding, we performed Y2H assays with CD domain mutations (D→N) that disrupt the two conserved Asp residues. MPK6 CD^D->N^ was unable to associate with both SCRM as well as the scrm-D KiDoK motif (Fig. 4a), indicating that MPK6 binds to SCRM using its conserved CD domain, possibly using a mechanism similar to that in mammalian MAPKs. These results were further supported by *in vitro* BLI measurements, which showed MPK6 CD^D->N^ failed to show appreciable affinity towards SCRM KiDoK (K_D_ value: N.F.) compared to the wild-type MPK6 protein (K_D_ value: 47±6 nM) (Fig. 4b). Taken together, the structure-function analyses of MPK6 and SCRM-KiDoK motif reveal a conserved mechanism of substrate binding between plant and animal MAPKs through the CD domain and the Docking motif of their diverse substrates. However, these results do not address how the KRAAM motif, that is shared only among SCRM orthologs, governs specific binding to MPK3/6.

**Fig. 4:**
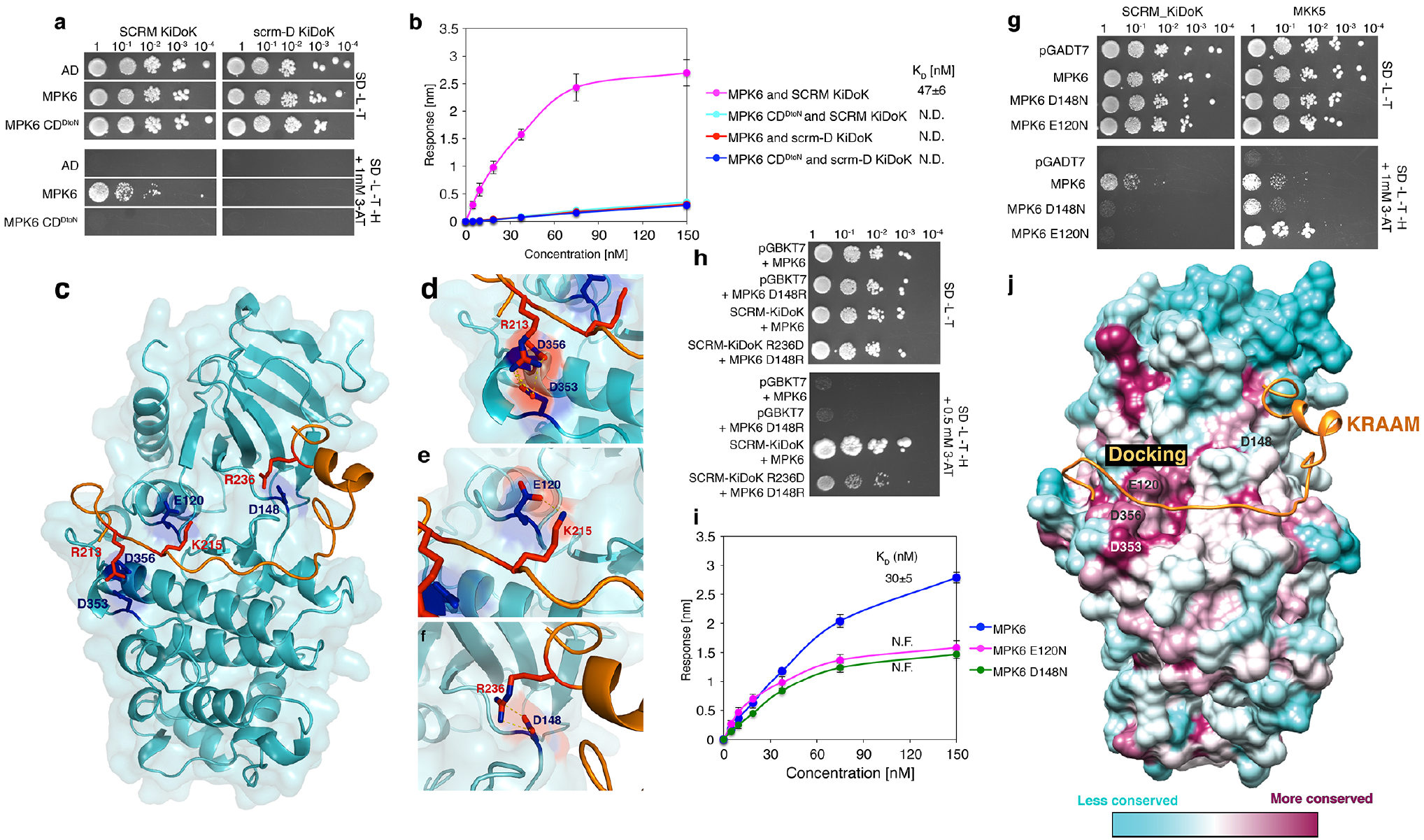
Structure-function analyses of the SCRM_KiDoK-MAPK interaction module. **a,** Y2H assays of SCRM-KiDoK and scrm-D KiDoK motif along with MPK6 and CD domain substitutions of MPK6. Bait: wild-type SCRM_KiDoK motif and the scrm-D KiDoK motif. Prey: MPK6, MPK6 D353N D356N (MPK6-CD^DtoN^). **b,** Quantitative analysis of interactions between the KiDoK motif of SCRM and scrm-D and GST MPK6/MPK6-CD^DtoN^ using BLI. Shown is the *in-vitro* binding response curve for recombinantly purified GST-MPK6/MPK6 CD^DtoN^ and biotinylated KiDoK motif peptide of SCRM/scrm-D at 6 different concentrations (150, 75, 37.5, 18.75, 9.375, 4.6875 nM). Values represent the mean and error bars the S.D. of three independent experiments performed. K_D_ values are listed next to the graph legend. **c,** Flexible docking of the *ab-initio* modeled SCRM peptide (orange) onto the crystal structure of MPK6 resolved in this study (cyan). The contact residues are shown in blue on MPK6 and red on SCRM KiDoK peptide. The non-terminal missing regions of the MPK6 structure (chain B) were modeled using Modeler and the resulting structure was used as an input for the Rosetta FlexPepDock program along with the 35-residue sequence of the SCRM KiDoK peptide. **d,** Zoomed-in view of R213 (red) on SCRM (orange) and D353/D356 (blue) on MPK6 (cyan). Polar contacts are represented by yellow dotted lines. Oxygen is shown in red and nitrogen is shown in blue. **e,** Zoomed-in view of K215 (red) on SCRM (orange) and E120 (blue) on the CD motif of MPK6 (cyan). Polar contacts are represented by yellow dotted lines. **f,** Zoomed-in view of R236 (red) on SCRM (orange) and D148 (blue) of MPK6 (cyan). Polar contacts are represented by yellow dotted lines. **g,** Y2H assays. Bait: SCRM KiDoK motif and MKK5. Prey: the activation domain alone (AD), MPK6, MPK6 D148N and MPK6 E120N. While the D148N and the E120N mutation in MPK6 dramatically reduces the interaction between SCRM KiDoK and MPK6, the mutation does not alter the interaction with other MPK6 substrates such as MKK5. **h,** Residue swap assays. Bait: DNA binding domain alone (pGBKT7), SCRM KiDoK motif, SCRM KiDoK R236D motif. Prey: MPK6 and MPK6 D148R. Swapping the residues of the D148 and R236 contact sites of MPK6 and SCRM still maintained interaction between the mutant variants of MPK6 and the SCRM KiDoK motif, albeit to a lesser extent compared to its wild-type counterparts. **i,** Quantitative analyses of the interactions between SCRM KiDoK motif and GST-MPK6, GST-MPK6 D148N, GST-MPK6 E120N using BLI. Shown is the *in-vitro* binding response curve for GST-MPK6, GST-MPK6 D148N and GST-MPK6 E120 with SCRM KiDoK peptide at 6 different concentrations (150, 75, 37.5, 18.75, 9.375, 4.6875 nM). The D148N as well as the E120N mutations in MPK6 dramatically lower the binding response as well as the K_D_ values for their interactions with the SCRM peptide. Values represent the mean and error bars the S.D. of three independent experiments performed. K_D_ values are listed on top of the response curves. **j,** Surface conservation mapping of MPK6 (cyan-maroon) with the SCRM peptide (orange). Surface coloring reflects sequence conservation between *A. thaliana* MPK6 and *H. sapiens* ERK1, ERK2, ERK5, p38α, p38β, p38γ, and p38δ. Maroon patches represent more conserved regions and cyan patches represent less conserved regions. MPK6 residues that bind to the SCRM Docking motif i.e. E120, D353, and D356 are significantly more conserved than the D148 contact site, which specifically binds with the SCRM KRAAM motif, indicating substrate specificity in MAPKs with respect to the functional outcomes they control.

### Deciphering the MPK6-SCRM binding interface

To understand the structural basis behind how the single residue substitution within the SCRM KRAAM motif abolishes association with MAPKs, we initially sought to resolve the crystal structure of the MPK6-SCRM_KiDoK complex, which was unfortunately recalcitrant to crystallization. As an alternative approach, we resorted to *ab-initio* modeling of the MPK6-SCRM protein-peptide complex and performed docking simulations of the SCRM_KiDoK peptide onto our AtMPK6 crystal structure (PDB ID: 6DTL) (Fig. 4c-f). Linear binding motifs that interact with the kinases are often devoid of tertiary structure and form a well-defined structure only in the presence of its binding partner ^28^. In light of this, flexible docking was performed wherein the SCRM_KiDoK peptide was allowed to fold near the CD domain and docking groove with multiple constraints imposed using information from previous studies 11,27,28 as well as experiments defined in this study (See Methods). The SCRM Docking motif was restrained to the D-motif binding site, which is composed of the CD domain and the hydrophobic docking groove ^11^, and the KRAAM motif was restrained to be in the vicinity of the MPK6 structure. The R213 residue of the SCRM Docking motif was restrained to remain near the negatively charged CD domain residues D353 and D356 (Fig. 4d). Comparison of previously determined ERK2-substrate peptide complexes (ERK2-pepMNK1, PDB ID: 2Y9Q and ERK2-pepRSK1, PDB ID: 3TEI)^27^ to our MPK6 crystal structure highlights possible interaction between the positively charged residue K215 of the SCRM Docking motif and E120 of MPK6 (Fig. 4e).

Next, we performed functional analyses to validate the two top-scoring final models. One model predicted MPK6 E163 and E164 as possible interaction sites with R236 of the KRAAM motif. This was rejected by the experimental validation, since site-directed mutations within these MPK residues to eliminate negative charges (E163N and E164N) showed no effects on SCRM interaction (Fig. S5). The other model predicted D148 of MPK6 as a potential interaction partner of R236 of the KRAAM motif, which was fully supported by further experimental verifications (See next section).

### Bipartite recruitment of MPK6 by SCRM using a conventional docking site and a SCRM-specific motif

The MPK6-SCRM interaction model predicts the bipartite binding mode of SCRM to MPK6: The Docking motif associating with the CD domain of MPK6; The KRAAM motif lies at the neck of MPK6 N- and C-terminal lobes with MPK6 D148 as a direct interaction site to SCRM R236, the key residue within the KRAAM motif (R236 is mutated to H in *scrm-D*) (Fig. 4). Importantly, the MPK6 D148 site is not conserved amongst mammalian MAP kinases (Fig. S5b). To investigate the significance of MPK6 D148 and SCRM R236 as a binding interface, we first performed site-directed mutagenesis. Substitution of MPK6 D148 to N dramatically compromised its interaction with the SCRM_KiDoK motif in Y2H as well as *in vitro* quantitative binding kinetics assays using BLI (Figs. 4g, I, S5a), indicating that the D148 is indeed critical for binding with SCRM through R236 within the KRAAM motif.

Next, to address whether the polar interactions between MPK6 D148 and the SCRM R236 are sufficient for MPK6-SCRM association, we swapped these residues to D148R and R236D, respectively. As shown in Fig. 4H, indeed, MPK6 D148R and SCRM R236D were still able to maintain interaction, albeit to a lesser extent than their wild-type counterparts. This suggests that these two D and R residues serve a critical interaction interface even when they are placed out of context from their native protein environments.

To gain further insight into the bipartite binding mode, we performed surface conservation mapping of AtMPK6-SCRM complex and *H. sapiens* ERK1, ERK2, ERK5, p38 *α*, p38 *β*, p38 *γ*, and p38 *δ*. While AtMPK6 residues binding the SCRM Docking motif (i.e. D353, D356 and E120) are significantly more conserved with mammalian MAPKs, the D148 contact site is far less conserved, thus reflecting the uniqueness of SCRM KRAAM motif binding to MPK6 (Figs. 4g, h, j). Finally, to decipher the specific role of MPK6 D148 while associating with SCRM KRAAM motif, we examined whether the MPK6 D148N substitution, which disrupts association with SCRM, has any effects on MPK6 for recruiting an upstream MAPKK, MKK5 ^29^. Indeed, MPK6 D148N retained interaction with MKK5 at a similar level to wild-type MPK6 (Fig. 4g). An additional E120N substitution also retained MKK5 association but compromised binding to SCRM_KiDoK (Figs. 4g, j), further emphasizing the intricate nature of MPK6-SCRM binding. Taken together, our structural and *ab initio* modeling of the MPK6-SCRM KiDoK complex and further experimental verifications elucidate a unique, bipartite binding mode of SCRM with MPK6: one that uses both conserved substrate binding sites similar to animal MAPKs, as well as contact sites that are highly unique to SCRM, the union of which results in specialized direction of cell-fate during stomata development.

### Binding of SCRM-KiDoK and MPK3/6 is critical role for SCRM phosphorylation and stability

To probe whether MPK3/6 can modify SCRM but spare scrm-D, we first performed an *in vitro* kinase assay with purified, recombinant MKK5^DD^, MPK3, MPK6, SCRM and scrm-D (see Methods). In the presence of MPK3 and MPK6 that is activated by a constitutively active upstream MAPKK (MKK5^DD^), we detected strong phosphorylation of GST-SCRM as previously reported in the context of freezing tolerance ^30, 31^. By constrast, GST-scrm-D is not phosphorylated by either MPK3 or MPK6 (Fig. 5a).

**Fig. 5:**
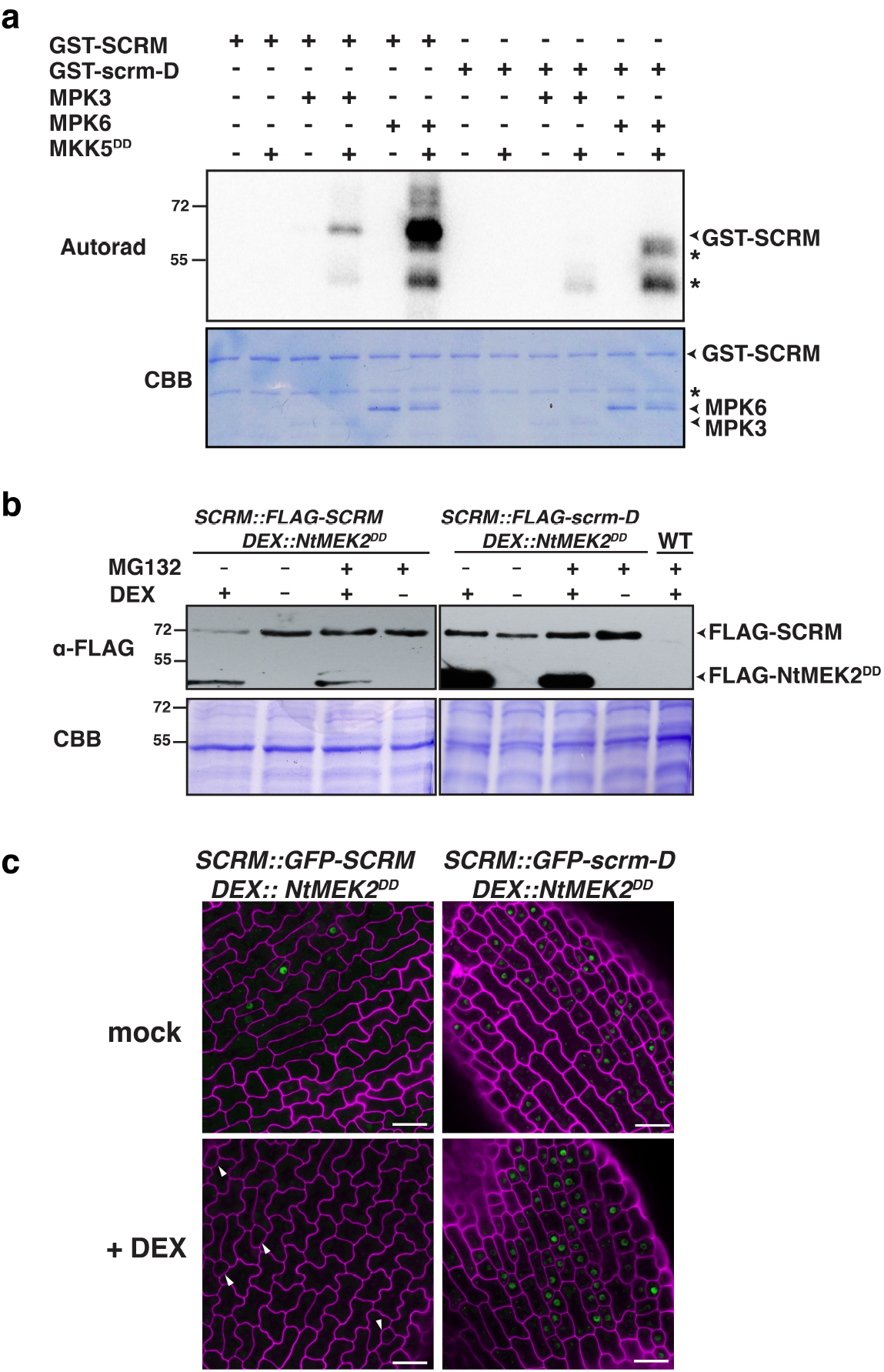
Direct MPK3/6 association is required for the phosphorylation and degradation of SCRM. **a,** *In vitro* phosphorylation assay of SCRM and scrm-D. Purified, recombinant SCRM, scrm-D, MKK5^DD^, MPK3 and MPK6 were subjected to *in vitro* phosphorylation followed by SDS-PAGE. Autoradiograph (top); Commassie Brilliant Blue (CBB: bottom). Asterisks: non-specific signals. **b,** Induced NtMEK2^DD^ overexpression triggers *in vivo* protein degradation of SCRM, but not scrm-D. Transgenic *Arabidopsis* seedlings carrying *DEX::FLAG-NtMEK2^DD^* with *SCRMpro::3xFLAG-SCRM* or with *SCRMpro::3x-FLAG-scrm-D* were grown in the presence of 0.5 μM DEX (+) or mock (−). At 4 days-post-germination (dpg), the seedlings were treated with the proteasomal inhibitor MG132 (+; 50 μM) or mock (−) for 6 hours. The total proteins were separated by SDS-PAGE (bottom; CBB) and subjected to Immunoblot with anti-FLAG antibody (top; *α*-FLAG). **c,** Induced NtMEK2^DD^ overexpression triggers loss of GFP-SCRM but not GFP-scrm-D. Shown are cotyledon abaxial epidermis from 1-day-old double transgenic *Arabidopsis* seedlings carrying inducible NtMEK2^DD^ (*DEX::FLAG-NtMEK2^DD^*) with *SCRMpro::GFP-SCRM*, or with *SCRMpro::GFP-scrm-D* grown in the presence of 0.05 μM DEX (+Dex) or ethanol (mock). Upon NtMEK2^DD^ induction, GFP-SCRM signals become undetectable throughout the entire epidermis, including those cells that would have been GMCs (white arrowheads), whereas the GFP-scrm-D signals still persist. Scale bars = 20 μm.

To unravel the *in vivo* consequence of the MPK3/6-mediated SCRM phosphorylation, we first generated double transgenic Arabidopsis seedlings expressing a functional, epitope-tagged SCRM as well as scrm-D driven by the endogenous promoter (*SCRMpro::FLAG-SCRM* and *SCRMpro::FLAG-scrm-D*) and an inducible constitutively-active MAPKK, *DEX::FLAG-NtMEK2^DD^*, which has been widely used to activate Arabidopsis MPK3/6 *in vivo* 10,32 (Fig. 5b). Upon DEX induction, the SCRM protein readily degrades, whereas upon treatment with the proteasome inhibitor MG132 in the presence of DEX induction, the protein accumulation level of SCRM is restored in the *SCRMpro::FLAG-SCRM* seedlings (Fig. 5b), indicating that SCRM is subjected to proteasome-mediated degradation upon induction of MAPK-mediated phosphorylation. In contrast, the scrm-D protein remained stable regardless of the NtMEK2^DD^ induction or proteasome inhibitor treatment (Fig. 5b).

Next, we examined the *in vivo* stability of SCRM and scrm-D proteins during seedling epidermal development using double transgenic seedlings expressing functional GFP-fused SCRM and scrm-D and *DEX::FLAG-NtMEK2^DD^* (Fig. 5c). While NtMEK2^DD^ induction triggered loss of GFP signals from the epidermis of *SCRMpro::GFP-SCRM*, including those cells that appeared to have entered the stomatal lineage (Fig. 5c, white arrowheads), the GFP signals from *SCRMpro::GFP-scrm-D* seedlings persist even after the activation of MAPKs *in vivo* (Fig. 5c). On the basis of these results, we conclude that the direct association of SCRM and MPK3/6 via the KiDoK motif is critical for subsequent phosphorylation and degradation of the SCRM protein, thereby specifying proper cell fate during stomatal differentiation (Fig. 6).

**Fig. 6:**
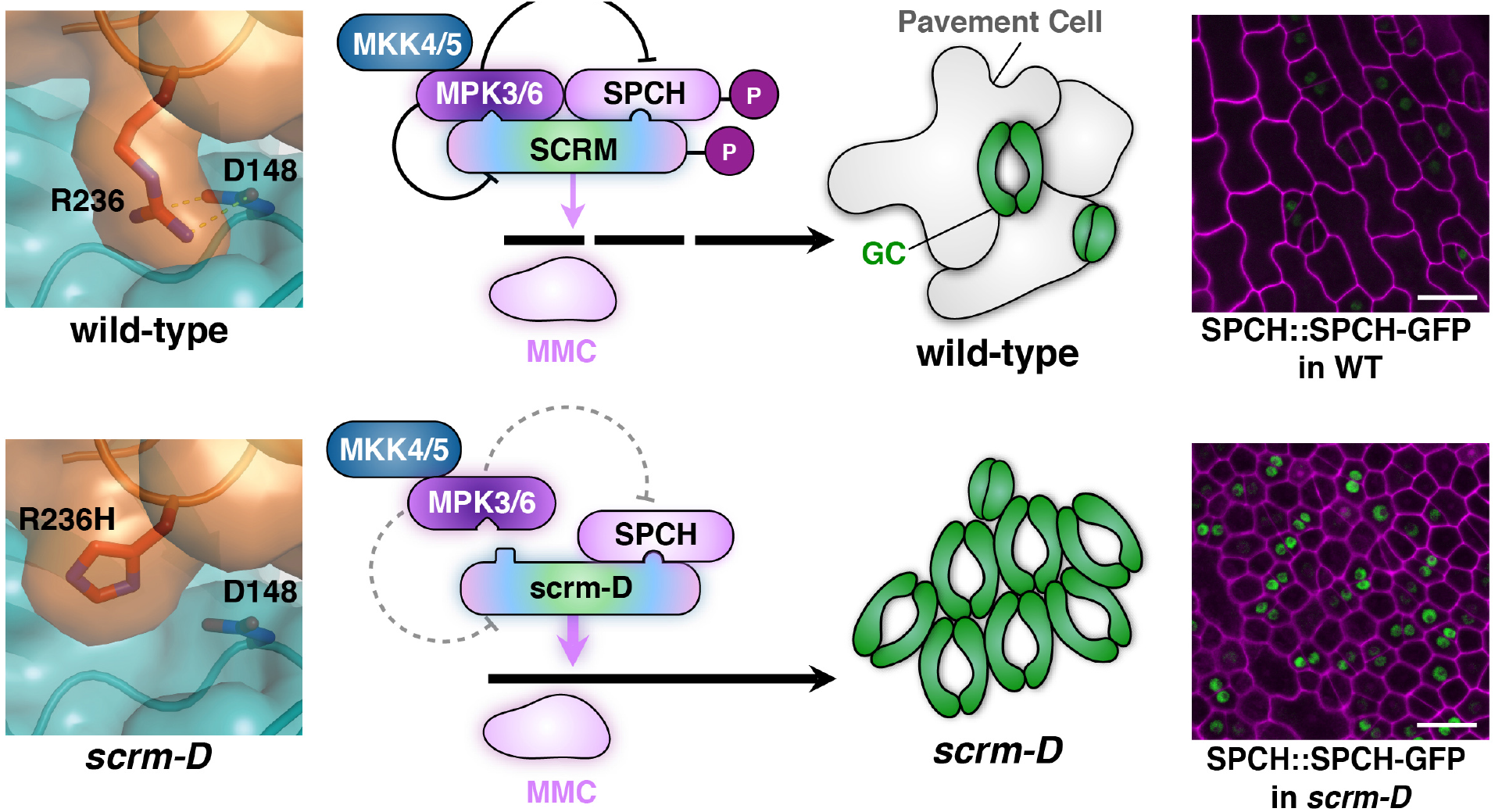
Mechanism enforcing the initiation of stomatal cell lineages via SCRM_KiDoK-MAPK interaction module. Shown is a schematic diagram depicting the molecular interaction mechanism between MPK3/6, SCRM, and SPCH for proper specification of stomatal cell-fate on the developing epidermis. In wild-type plants, The KiDoK motif of SCRM associates with MPK3/6 through its bipartite Docking and KRAAM motifs. This association of MPK3/6 and SCRM triggers subsequent phosphorylation and degradation of the SCRM protein by an unknown proteosomal pathway. Through heterodimerization with SPCH, SCRM brings SPCH to a molecular proximity with MPK3/6, thereby allowing the phosphorylation-mediated downregulation of SPCH to inhibit stomatal cell fate. In *scrm-D*, since the scrm-D_KiDoK motif (K<u>H</u>AAM) is unable to associate with MPK3/6, the scrm-D protein cannot recruit MPK3/6 to interact with and modulate the stability of itself and SPCH, resulting in a highly-stabilized SPCH protein in the *scrm-D* background (Right). The absence of proper downregulation of SCRM and SPCH confers constitutive stomatal differentiation in the entire epidermis.

## Discussion

Our study elucidates, at the atomic resolution, the mechanism by which SCRM integrates upstream MAPK cascade repressors and the downstream transcription factor SPCH to enforce the initiation of stomatal cell lineages on the developing plant epidermis (Fig. 6). Both the Kinase Docking and KRAAM motifs of SCRM are necessary for MPK6 binding, whereas the former motif appears dispensable for association with MPK3 (Fig. 3). The sequence conservation of the SCRM_KiDoK motif amongst SCRM orthologs (Fig. S2a) suggests their nuanced regulation by MAPKs. The high sequence conservation of the KRAAM motif within SCRM orthologs from both vascular and non-vascular plants correlates with their tight binding with AtMPK3, whereas the relatively low sequence conservation of the Kinase Docking motif correlates with their disparate binding with AtMPK6 (Figs. 2d, e). This is most evident in *P. patens* SCRM (PpICE1), which has a highly conserved KRAAM (KRAAS) motif but a poorly conserved Kinase Docking motif: PpICE1 associates strongly with AtMPK3 but very poorly with AtMPK6. These findings imply an elaboration of a bipartite binding mode of SCRM with MAPKs during the evolution and diversification of land plants. The cryptic functional non-redundancies between MPK3 and MPK6, unraveled through mutational analysis of the KiDoK motif (Fig. 3d) further hint at the hidden, unique properties associated with these two MAPKs. Among the SCRM orthologs, BdSCRM2 lacks the entire KiDoK region altogether, even though it can substitute for BdICE1 to produce normal stomata in *Brachypodium* ^33^. As expected from the amino-acid sequence, BdSCRM2 does not interact with AtMPK3/6 (Fig. S2c), suggesting that BdSCRM2 is not under direct regulation by the MAPK cascade. Therefore, grass species may have not only re-wired the core stomatal bHLH TFs ^33^, but also re-shaped the architecture of the repressive signaling pathway feeding into the stomatal differentiation program.

The inability of the scrm-D protein to associate with MPK3/6 prevents its phosphorylation and subsequent degradation (Figs. 5, 6). Thus, the KiDoK motif of SCRM essentially functions as a degron, controlling the stability of the SCRM protein. Over the years, a number of components that facilitate the degradation of SCRM in diverse biological contexts have been identified. For example, during freezing acclimation, an E3 ligase, HOS1 and SIZ1 mediate the ubiquitination and SUMOylation of SCRM/ICE1 and subsequent degradation ^34, 35^. Additionally, the OST1 kinase enhances SCRM/ICE1 stability by suppressing HOS1-mediated degradation of SCRM/ICE1 in the cold ^36^. Recently, another E3 ligase, COP1, has been shown to suppress stomatal development in the dark by ubiquitinating and degrading SCRM ^37^. This COP1-mediated SCRM degradation can be abrogated by light. As such, identification of the E3 ligase that regulates the SPCH·SCRM bHLH heterodimer module during normal stomatal development is an important future direction.

SCRM controls developmental and environmental processes outside of stomatal differentiation likely via associating with specialized bHLHs ^38, 39^. For instance, SCRM forms a heterodimer with ZHOUPI, a specialized bHLH protein involved in endosperm/seed development ^40^. Hence, our discovery that SCRM acts as a scaffold to recruit MPK3/6 to its partner bHLH, SPCH, could be a universal mechanism for MPK3/6-mediated regulation of transcriptional reprogramming in diverse contexts. Recent studies have shown that phosphorylation of SCRM/ICE1 by MPK3/6 is critical for plant freezing acclimation ^30, 31^. It is not known whether SCRM/ICE1 forms heterodimers with specialized bHLHs and regulates their phosphorylation as a scaffold for cold tolerance pathway.

The proposed bipartite binding mode of the SCRM KiDoK motif with MPK6 explains the complexity in binding specificity between MPK3/6 and SCRM. While mammalian MAPKs are known to interact with their substrates using a conserved Kinase Interaction Motif (KIM) [KR]X_2–6_[LI]X[LI] ^41^, this type of motif has been identified in plants only in APC21, an Arabidopsis type 2C Ser/Thr phosphatase that negatively regulates MPK4 and MPK6 and modulates innate immunity, jasmonic acid and ethylene levels ^42^. It seems rather unlikely that diverse MAPK substrates use similar KIMs to interact with MAPKs that are known to simultaneously influence a variety of developmental programs.

The highly conserved KRAAM motif is found only in SCRM/SCRM2 and its plant orthologs, reiterating the notion that distinct binding motifs in MAPK substrates are used to elicit unique developmental responses. Furthermore, it is also known that putative kinase interaction motifs in MAPK substrates rarely overlap with other highly conserved domains (e.g. bHLH domain), indicating that diverse MAPK substrates might not compete for the same binding site on MAPKs to specify a unique developmental outcome ^1^. From our structural and functional analyses of the MPK6-SCRM-KiDoK complex, the simultaneous association of the Kinase Docking and the KRAAM motifs of SCRM with MPK6 is required to dictate cell-fate specification during stomatal development. While it is evident from the surface conservation mapping of AtMPK6 and mammalian MAPK orthologs that the D148 residue of AtMPK6 is not as conserved with mammalian MAPK orthologs (Fig. 4j), it is perfectly conserved among Arabidopsis MAPKs (Fig. S5d). Conversely, a sequence conservation analysis among CD domain residues of all AtMAPK family members revealed the contact sites to be highly diverse (Fig. S5e). These findings explain why the bipartite binding mode is required for SCRM to uniquely interact with MPK6, but not with other MPKs like MPK4 (Fig. S2b), to specify cell-fate during stomatal development.

The scaffolding action of SCRM, as discovered through our structural and functional analyses, has shed light on its post-translational regulation during stomata development. While the wild-type SCRM enables MAPK to interact with SPCH and consequently regulate its activity through phosphorylation, scrm-D can no longer serve as a scaffold to facilitate this interaction due to its R236H substitution, which abolishes interaction with D148 of MPK6 (Fig. 6). A direct ramification is the enhanced stability of the SPCH protein in the *scrm-D* background, conferring constitutive stomatal differentiation (Fig. 6). The initiation of stomatal cell lineage is controlled by multiple signals, including the MAPK cascade, the brassinosteroid-mediated BIN2 kinase as well the ABA signaling pathway ^43-45^. By serving as a direct scaffolding partner for SPCH and MPK3/6, SCRM bridges the gap in our understanding of how multiple opposing signaling cues get integrated simultaneously into the highly regulated stomatal differentiation pathway to seamlessly deliver systematic cell-state transitions during plant development.

## Methods

### Plant materials and growth conditions

All *Arabidopsis* plants used in this study are in the Columbia (Col) background. *scrm, scrm2, scrm-D, SCRMpro::GFP-SCRM, SCRMpro::GFP-scrm-D, mpk3, mpk6 and MEK2DD* have been reported previously ^10, 21^. Transgenic plants expressing the SCRM-KRAAM motif substitutions (*SCRMpro::SCRM-KAAAM, SCRMpro::SCRM-ARAAM, SCRMpro::SCRM-AHAAM, SCRMpro::SCRM-AAAAM*) as well as the SCRM docking motif deletions (*SCRMpro::3XFLAG-SCRM-ΔKinase Docking and SCRMpro::3XFLAG-scrm-D-ΔKinase Docking*) in *scrm* and *scrm scrm2* backgrounds were generated in this study. Plants were grown at 21°C with long-day light cycle (16 h light/8 h dark). The primers used for genotyping and construct generation are listed in Table S3.

### Plasmid construction and generation of transgenic plants

For the compete list of plasmids generated in this study, see Table S3. To generate MPK6 ΔN terminus for recombinant protein expression, the CDS sequence of MPK6 (85-1188bp) was amplified using Phusion Polymerase and cloned into pGEX4T-1 using the BamHI and NotI restriction sites. For Y2H assays, the CDS sequence of the gene of interest was fused to either the DNA binding domain of vector pGBKT7 or the activation domain of vector pGADT7 using restriction sites EcoR1 and BamHI. For site-directed mutagenesis, complementary primers with corresponding mutation sites were designed for PCR amplification of the template plasmid using PrimeStar Polymerase. The PCR products were then treated with DpnI to digest the parental methylated DNA. The digested PCR products were purified using Qiagen PCR purification kit and then transformed into DH5α chemically competent cells. The mutated transformants were confirmed by sequencing. For BiFC assays, Split YFP constructs were generated by cloning the gene of interest into either the 35S::pSPYNE-GW vector ^24^, which contains of the N-terminal of EYFP protein (YFPn -174 amino acid) or 35S::pSPYCE-GW which contains the C-terminal of EYFP protein (YFPc -64–amino acid) using LR Gateway Recombination cloning methods ^24^. The constructs were then transformed into the Agrobacterium strain GV3101, and co-infiltrated along with the silencing suppressor plasmid p19 into *Nicotiana Benthamiana* leaves. For generating transgenic plants, constructs were transformed into the Agrobacterium strain GV3101, and transgenic plants were subsequently generated via floral dipping method ^46^. At least 6 T1 lines were phenotypically characterized and the lines showing the strongest phenotype were used in further studies.

### Yeast two-hybrid screen

Yeast two-hybrid (Y2H) screening was performed by Hybrigenics Services (Paris, France) (http://www.hybrigenics-services.com). The coding sequences for *Arabidopsis thaliana* SCRM/ICE1 (aa 207-297, <u>NM_113586.4</u>) was PCR-amplified and cloned into pB27 as a C-terminal fusion to LexA (orientation LexA-bait). The construct was checked by sequencing the entire insert and used as a bait to screen a random-primed Arabidopsis thaliana 1-week-old seedlings cDNA library constructed into pP6. pB27 and pP6 are derived from the original pBTM116 ^47^ and pGADGH ^48^ plasmids, respectively. 158 million interactions were screened using a mating approach with YHGX13 (Y187 ade2-101::loxP-kanMX-loxP, mat α) and L40 Gal4 (mat a) yeast strains as previously described ^49^. 50 Positive clones were selected on a medium lacking tryptophan, leucine and histidine. The prey fragments of the positive clones were amplified by PCR and sequenced at their 5’ and 3’ junctions. The resulting sequences were used to identify the corresponding interacting proteins in the GenBank database (NCBI) using a fully automated procedure. A confidence score (PBS, for Predicted Biological Score) was attributed to each interaction as previously described ^50^. See Table S1 for the full list.

### Yeast two-hybrid assay

Bait and prey constructs were co-transformed into the yeast strain AH109 using the yeast transformation kit (Frozen-EZ Yeast Transformation II Kit™ - Zymo Research). Y2H assays were performed using the Matchmaker 3 system (Clontech). The resulting transformants with appropriate positive and negative controls were spotted on SD (-Leu/-Trp) plates to check for growth in the absence of selection. The transformants were then spotted on SD (-Trp -Leu -His) selection media containing 0.5/1mM 3-Amino-1,2,4-triazole (3-AT; Sigma, A8056). The positive interactors were then scored based on the stringency of the selection.

### Bimolecular fluorescent complementation (BiFC) assay

BiFC assays were carried out as described previously^24^ with minor modifications. Split YFP constructs were generated for SCRM, scrm-D, MPK3, MPK6 and SPCH by cloning them into either the 35S::pSPYNE or the 35S::pSPYCE Gateway Recombination vectors ^24^. The constructs were then transformed into the Agrobacterium *tumefaciens* strain *GV3101*. 3ml LB media was inoculated with the agrobacterium transformants and incubated overnight with gentle shaking at 28°C. Next morning, cultures were spun down at 4500 rpm for 10 minutes and resuspended in infiltration buffer (10 mM MgCl2, 10 mM MES (pH 5.6) and 150 μM acetosyringone). Bacterial culture densities were adjusted to a final OD_600_ of 1.0, and the cell-suspensions were incubated at room temperature for 4 h prior to infiltration. Equal volumes of cultures carrying the corresponding complementary pair of BiFC constructs (YFPn and YFPc) along with silencing suppressor plasmid - p19 (a gift from Professor Sir David Baulcombe) ^51^ were then co-infiltrated into 3-4 week old N.*benthamiana* leaves. The infiltrated leaves were imaged using confocal microscopy two days post infiltration.

### Confocal microscopy

Confocal imaging for Arabidopsis seedlings was done using the Zeiss LSM700 Confocal microscope. Cell peripheries were visualized by staining with propidium iodide (Molecular Probes) and by using the following settings: excitation 619 nm, emission 642 nm. Excitation at 488 nm and emission at 500-515 nm was used for visualizing GFP signals. Confocal Imaging for *N.Benthamiana* leaves was done using the Leica SP5x confocal microscope simultaneously capturing YFP (using the White Light Laser-excitation at 518 nm and emission at 540 nm for EYFP) and bright field channels. The confocal images were linearly adjusted uniformly for brightness/contrast using Photoshop CS6 (Adobe).

### Proteasome inhibitor treatment

*Arabidopsis* seedlings carrying dexamethasone-inducible NtMEK2^DD^ (*DEX::FLAG-NtMEK2^DD^*) with *SCRMpro::GFP-SCRM* and *SCRMpro::GFP-scrm-D* were grown in sterile petri dishes containing water for 4 days, following which 50μM MG132 (equal volume of ethanol for the mock-control) and/or 0.5μM DEX(equal volume of ethanol for the mock-control) were added and incubated for 6 hours, following which the tissue was harvested and flash frozen in liquid nitrogen. Total protein was then extracted using extraction buffer containing 50mM Hepes pH7.5, 150mM NaCl, 1mM EDTA, 1% Triton X-100, 0.1% sodium deoxycholate, 0.1% SDS, 100μM PMSF and 1:100 Protease Inhibitor Cocktail. 30 μg of total protein was loaded on an 8% SDS-PAGE gel, following which the proteins were transferred from gel to membrane and probed using Anti-FLAG antibody (Sigma).

### Recombinant protein expression and purification

A. *thaliana* MPK6 encompassing residues 29-305 (MPK6ΔNt) was cloned and expressed in E. coli with an N-terminal GST tag and a Thrombin cleavage sequence. Initial purification from cell lysate was performed using a glutathione agarose column. After loading and washing the column the GST tag was removed using an on-column Thrombin cleavage reaction to elute MPK6ΔNt. Further purification was performed with anion exchange and gel filtration columns. Final protein elution buffer consisted of 20mM Tris pH 8.0, 200mM NaCl, and 5mM DTT.

### Crystallization conditions and data collection

MPK6ΔNt crystals were grown at 4°C using hanging drop vapor diffusion. 1.5μl of protein was mixed with 1.5μl mother liquor (0.1M sodium citrate pH 6.5, 16% PEG 8000) and suspended from a glass slide. Crystals were harvested, incubated in 25% glycerol as a cryoprotectant, and flash frozen in liquid nitrogen. Diffraction data was collected remotely from the Advanced Light Source at Lawrence Berkeley National Laboratory on the BL8.2.1 beamline, and diffraction data were indexed, integrated and scaled with HKL2000.

### Crystal structure determination and refinement

The MPK6ΔNt structure was solved by molecular replacement using the Phaser-MR program in the Phenix software suite v1.10.1-2155 and PDB 5ci6 as a search model. An initial MPK6ΔNt structure was created with AutoBuild and subsequently refined over successive cycles with Coot v0.8.6.1 and phenix.refine. All structure figures were created using PyMOL (https://pymol.org/). See Table S2 for statistics from crystallographic analysis.

### Modeling and Docking of SCRM peptide to MPK6 crystal structure

The N-terminal missing regions of the MPK6 structure (chain B) were modeled using Modeler ^52^. This modeled MPK6 structure was used as input for Rosetta flexpepdock program ^53^ along with the 35 residue long amino-acid sequence of the SCRM peptide for ab-initio flexible docking. Initial structure of peptide sequence was modeled using the ab-initio modeling protocol of Rosetta Software Suite^54, 55^. For the initial docking model, constraints were imposed to keep the C*α* atoms of the R213 and K215 within 10 Å of C*α* atoms of the CD domain conserved residues D353 and D356. Further, residues L217, K218, E221 and L223 of SCRM were constrained to be within 10 Å of the C*α* atoms of the residues Y168, N200, H165, L197, I154 and L161 of MPK6. These constraints were determined from previous studies on MAP Kinase-peptide interactions^11, 27, 28^. Finally, the C*α* atoms of the KRAAM motif (residues 235-239) were restrained to remain within 10 Å of the MPK6 molecule. For the final models, side chain constraints were used to restrain the R213 near D353 and D356. Side chain of K215 was restrained to interact with E120 of SCRM. Additionally, side chain of R236 was restrained to interact with D148 of MPK6. The rest of the constraints were similar to the initial run with a shorter distance cutoff of 8Å.

### Surface Conservation Mapping

FASTA alignments of *A. thaliana* MPK6 and *H. sapiens* ERK1, ERK2, ERK5, p38, p38, p38, and p38 were generated with Qiagen Bioinformatics CLC Sequence Viewer v8.0.0. To map these onto our MPK6 structure, the alignments were exported into UCSF Chimera v1.11, developed by the Resource for Biocomputing, Visualization, and Informatics at the University of California, San Francisco, with support from NIH P41-GM103311. The Chimera MSMS package and POV-Ray v3.6 were used to model solvent-excluded molecular surfaces and generate raytraced images.

### Protein extraction and immunoblotting

Agroinfiltrated *N.Benthamiana* leaves expressing the desired constructs along with p19 were harvested 2 days post infiltration and ground in liquid nitrogen. The proteins were extracted using the following extraction buffer: 100 mM Tris-HCl pH 8.0, 150 mM NaCl, 10% Glycerol, 1% Triton X-100, 10mM Sodium Fluoride, 1mM Sodium Orthovanadate, 1mM EDTA, 1 mM PMSF and 1:100 Protease Inhibitor Cocktail. The frozen ground tissue was ground again in extraction buffer and the total extract centrifuged at 16,000g at 4°C for 20 min. The supernatant was transferred to a new tube. The protein concentration was measured using the Bradford Reagent (Bio-Rad) and final protein concentration of each sample was adjusted 20ug/lane for loading on an SDS PAGE Gel. The proteins were separated by SDS-PAGE (10% acrylamide gel) and transferred to a nitrocellulose transfer membrane (Immunobilon – EMD Millipore) using a semi-dry transfer set up. The samples were transferred at 120mA for 2 hours. The membranes were then blocked in 1x TBST buffer containing 5% non-fat dry milk powder following which they were incubated primary antibody (in 5% milk, 1x TBST) overnight. The membranes were washed 3 times using 1x TBST and then incubated with secondary antibody (in 5% milk, 1x TBST) at room temperature for 2 hours. The membranes were washed again 3 times using 1x TBST following which the bands were detected using Supersignal West Femto High Sensitivity Substrate (1:1).

### Recombinant protein expression and *in vitro* kinase assay

For *in vitro* protein expression, the constructs of pGEX-4T-1-SCRM, pGEX-4T-1-scrm-D, pET-28a-MPK3, pET-28a-MPK6, and pET-28a-MKK5^DD^ were transformed to *Escherichia coli* strain BL21. For each transformant, a single clone was selected and incubated in 3 ml LB liquid medium. The overnight-incubated *E. coli* suspensions were transferred to 300 ml LB medium and incubated at 37°C for ∼3 h until the OD_600_ reaching to 0.4-0.6. The isopropyl β-D-1-thiogalactopyranoside (0.5 μM of final concentration) was added to the cultures and the strains were incubated at 25°C for additional 5 h. SCRM and scrm-D proteins were purified by using glutathione agarose resin, and MPK3, MPK6, and MKK5^DD^ proteins were purified by using Ni-NTA agarose according to the manufacturer’s protocol. For *in vitro* kinase assay, two steps of reactions were conducted. Firstly, MPK3 or MPK6 protein alone or together with MKK5^DD^ were incubated in 30 μL of reaction buffer: 25 mM Tris-HCl (pH 7.5), 12 mM MgCl_2_, 1 mM DTT, and 50 μM ATP. The mixtures were kept at room temperature for 30 min. In the second step, 5 μL solution from each of the first reactions was incubated with SCRM/scrm-D in 25 μL of reaction buffer: 25 mM Tris-HCl (pH 7.5), 12 mM MgCl_2_, 1 mM DTT, 50 μM ATP, and 0.1 mCi [γ-^32^P]ATP. The reactions were processed for 30 min at room temperature before being stopped by adding SDS loading buffer. Proteins were separated by SDS-PAGE using a 10% (w/v) acrylamide gel, and the phosphorylated proteins were visualized by autoradiography.

### BioLayer interferometry (BLI)

Binding affinity of the SCRM KiDoK full-length peptide and the scrm-D KiDoK ful-length peptide with GST-tagged MPK3 and MPK6 were measured using the Octet Red 96 (ForteBio, Pall Life Sciences) following the manufacturer’s protocols. The optical probes coated with Streptavidin were first loaded with 100 nM biotinylated peptide (ligand) prior to kinetic binding analyses. The experiment was performed in black 96 well plates maintained at 30°C. Each well was loaded with 200μl reaction volume for the experiment. The binding buffer used in these experiments contained 1x PBS supplemented with 0.1% BSA. The concentrations of the GST-MPK3 as the analyte in the binding buffer were 1000, 333.33, 111.11, 37.03, 12.34 and 4.11nM. The concentrations of the GST-MPK6 as the analyte in the binding buffer were 150, 75, 37.5, 18.75, 9.375 and 4.6875nM. All preformed complexes remained stable as suggested by the constant signal during the post-loading washing step. There was no binding of the analyte to the unloaded probes as noted from the control wells. Binding kinetics to all 6 concentrations of analyte were measured simultaneously using default parameters on the instrument. The data was analyzed by the Octet data analysis software. The association and dissociation curves were fit with the1:1 homogeneous ligand model. The Kobs values were used to calculate the dissociation constant, *KD*, with steady state analysis of the direct binding. See Figure S6 for raw sensogram data.

### Data and software availability

The PDB accession number for MPK6Δnt structure reported in this paper is 6DTL.

## Acknowledgments

We thank Prof. Shuqun Zhang for his generous gifts of Arabidopsis *mpk3, mpk6*, and inducible *NtMEK2^DD^*overexpression line, and Prof. Juan Dong for thoughtful discussion. We thank Dr. Thomas Hinds for helping set up the *in vitro* interaction assays using the Octet system. This work was supported by the US National Science Foundation (MCB-0855659) and the Gordon and Betty Moore Foundation (GBMF-3035) to K.U.T., and Grants in Aid for Scientific Research on Innovative Areas, JSPS (26119006 & 15K21711) to F.T. N.Z. and K.U.T. are HHMI Investigators.

## Author contributions

Conceived the project, K.U.T.; Conceptualization, A.P., K.U.T.; Designed Experiments, A.P., A.L.R., N.Z., K.U.T.; Performed Experiments, A.P., J.R., C.Z., A.L.R., A.K.H., X.T.; Performed structural analysis and modeling; J.R., A.S., F.T., N.Z.; Formal Analysis, A.P., J.R., A.S., A.L.R., N.Z., K.U.T.; Visualization, A.P., J.R., A.S., K.U.T.; Writing-Original Draft, A.P., K.U.T; Writing-Review & Editing, A.P., J.R., A.S., C.Z., A.L.R., A.K.H, F.T., N.Z., K.U.T; Supervision, N.Z., K.U.T., Project Administration, K.U.T.; Funding Acquisition, J.K.Z., F.T., N.Z., K.U.T.

---

**Fig. S1:**
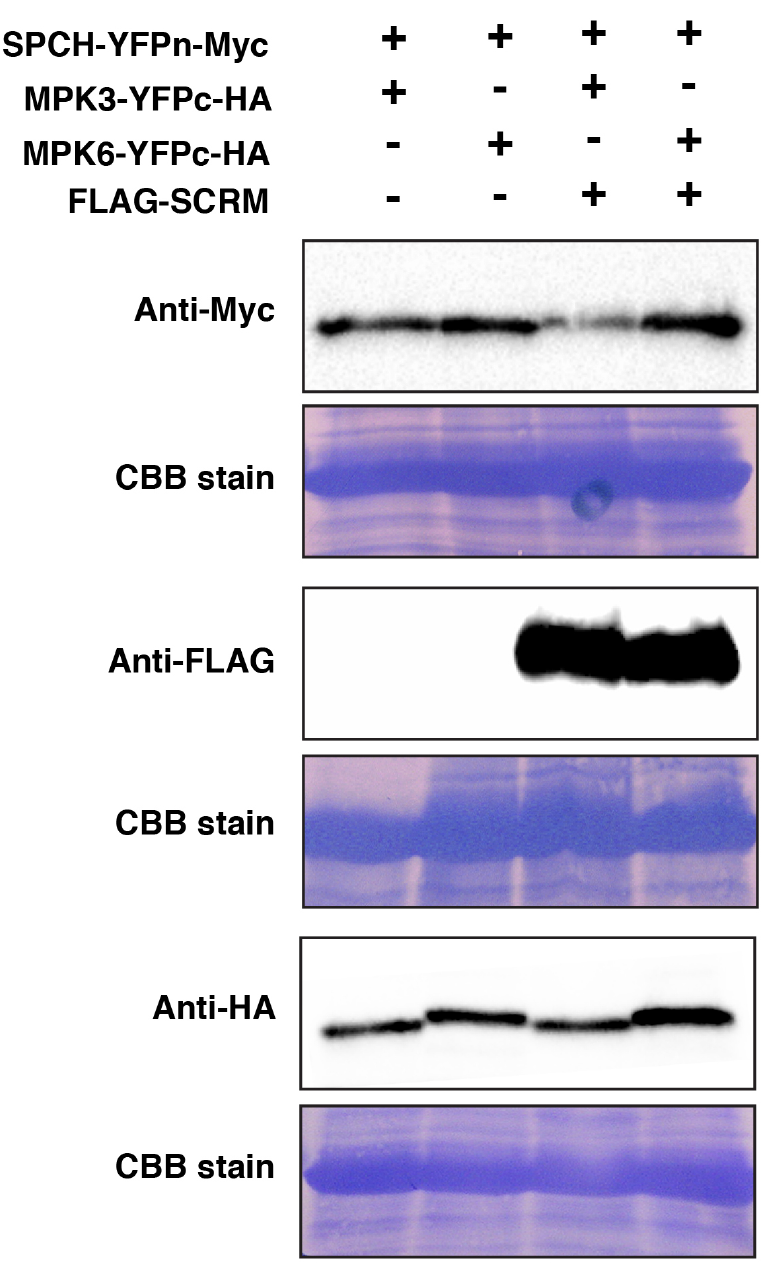
SPCH protein is expressed in the absence and presence of SCRM. (Related to Fig. 1). Shown are immunoblots for SPCH, SCRM and MPK3/6 protein expression detected using Anti-Myc, Anti-FLAG and Anti-HA antibodies respectively from 3-week old *N. benthamiana* leaves agroinfiltrated using pairwise combinations of SPCH-YFPn-Myc and MPK3-YFPc-HA, MPK6-YFPc-HA along with 35S:: FLAG-SCRM. SPCH protein accumulates both in the absence and in the presence of the SCRM protein, indicating that the absence of interaction between SPCH and MPK3/6 is not due to the SPCH protein not being expressed in the absence of SCRM. CBB staining for each immunoblot is presented below the same.

**Fig. S2:**
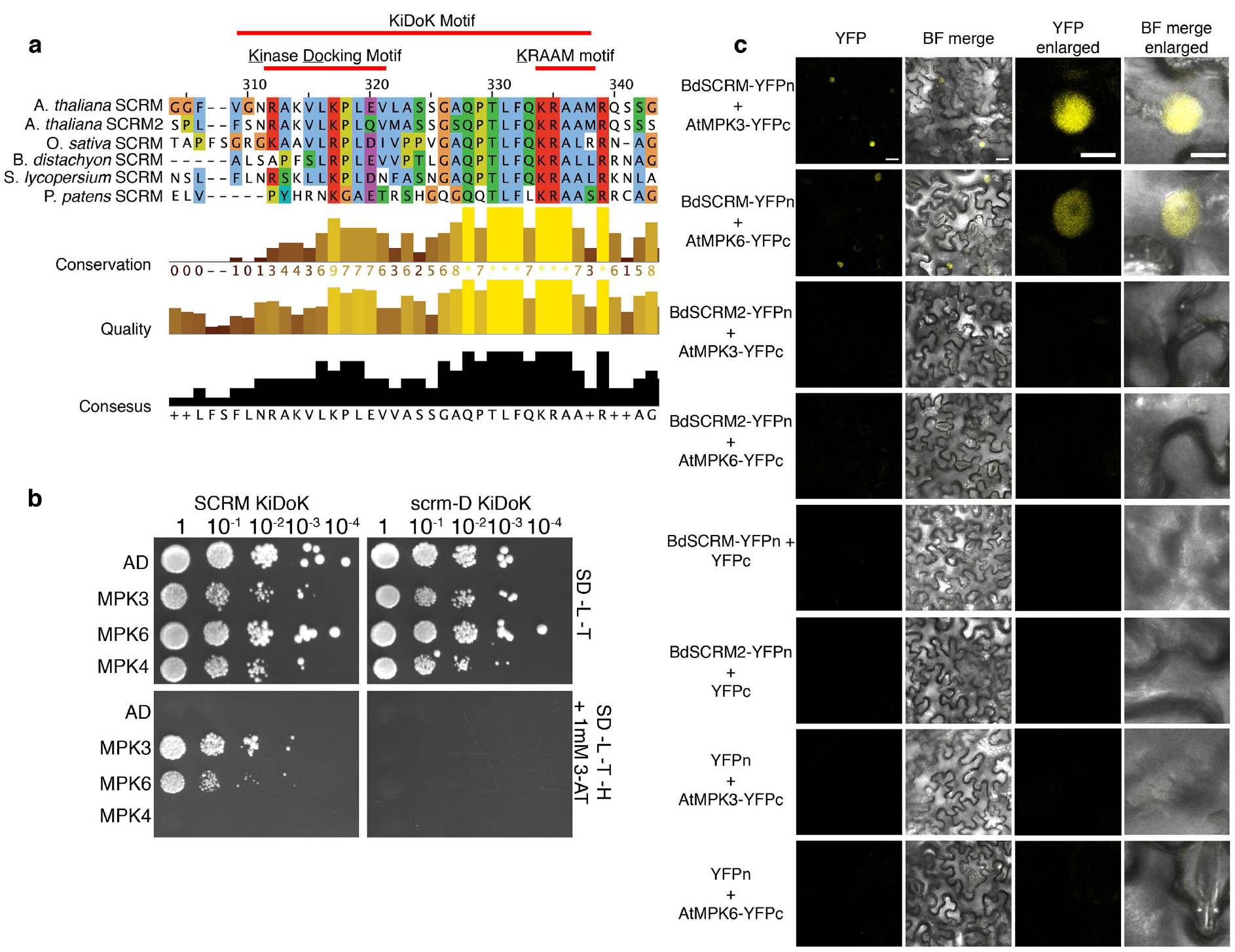
Sequence alignment and interaction between SCRM_KiDoK-MAPK in homologs/orthologs of AtMAPK and SCRM respectively. (Related to Fig. 2) **a,** Sequence alignment of the SCRM KiDoK motif in vascular as well as non-vascular plant orthologs of SCRM created using JALVIEW. The KiDoK motif, which includes the Kinase Docking motif and the KRAAM motif, are highlighted using the red line above the motif. **b,** Yeast two-hybrid (Y2H) assays of SCRM-KiDoK and scrm-D KiDoK motifs with AtMPK3, AtMPK6 and AtMPK4. Bait constructs containing the wild-type KiDoK motif (SCRM_KiDoK) and scrm-D version of the KiDoK motif (scrm-D_KiDoK) were tested in pairwise combinations with prey constructs containing the activation domain alone as well the activation domain fused to MPK3, MPK6 and MPK4. While MPK3 and MPK6 interact with SCRM KiDoK motif and not the scrm-D KiDoK motif, MPK4 does not interact with both. **c,** Bimolecular Fluorescent Complementation (BiFC) assays. Shown are 3-week old *N. benthamiana* leaves agroinfiltrated with pairwise combinations of BdSCRM-YFPn and BdSCRM2-YFPn along with AtMPK3-YFPc and AtMPK6-YFPc. Scale bar = 25 μm. Right two panels are magnified images of a representative nucleus. Scale bar = 10 μm. While BdSCRM associates with both AtMPK3 and AtMPK6 in the nucleus *in planta*, BdSCRM2 that lacks the KiDoK motif does not.

**Fig. S3:**
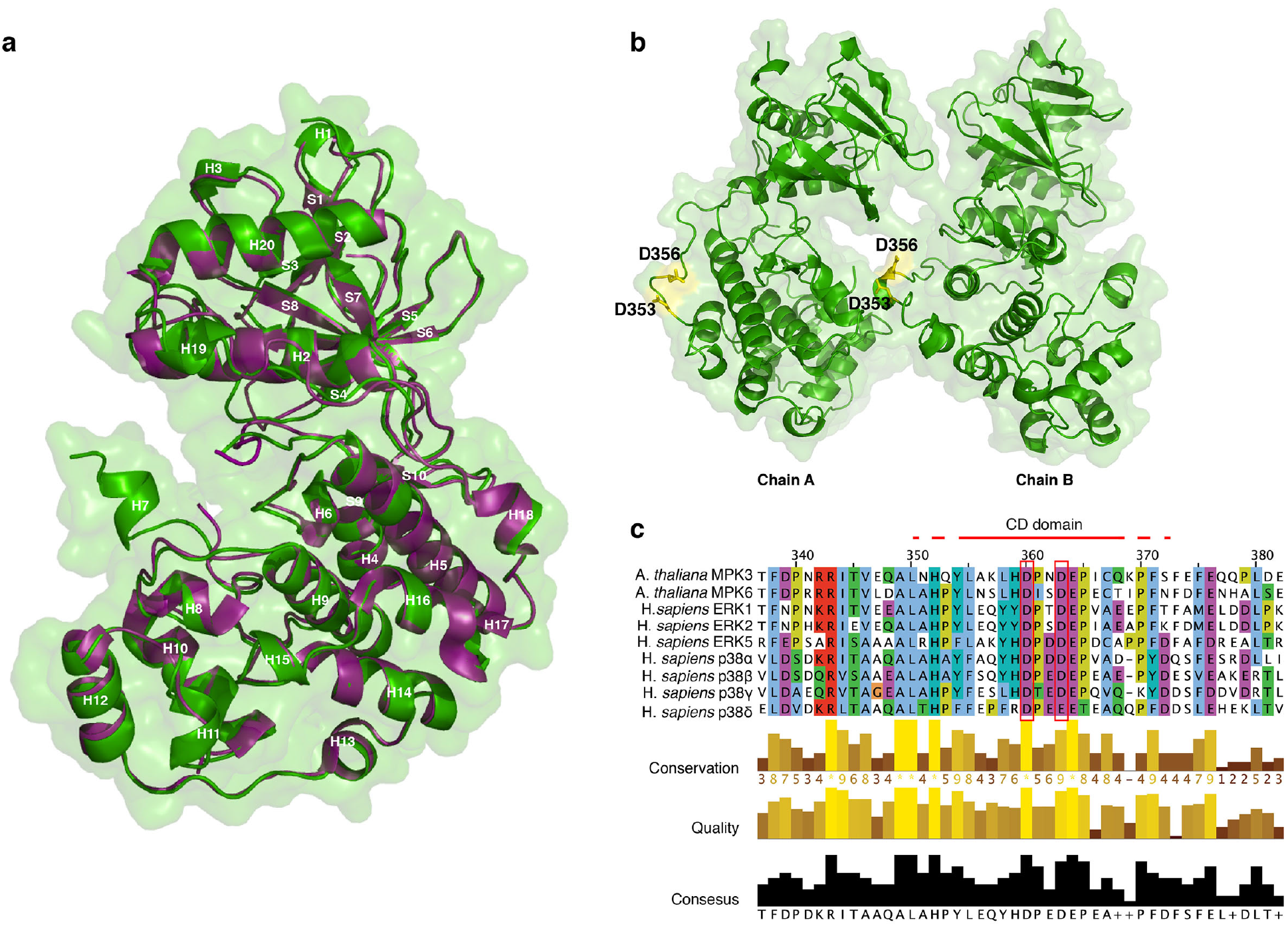
Crystal structure of AtMPK6, CD domain of AtMPK6 and sequence conservation within the CD domain of MAPKs. (Related to Fig. 4) **a,** Ribbon diagram superposition of the two MPK6 structures (green, this study PDB ID: 6DTL and magenta, PDB ID: 5CI6). MPK6 is composed of N-terminal and C-terminal lobes flanking a central ATP-binding region. Helices (H1-H20) are distributed between the N- and C-terminal lobes. Strands (S1-S10) are confined to the N-terminal lobe. The 6DTL includes an additional helical region, H7, comprising the phosphorylation loop proximate to the ATP-binding region. **b,** Ribbon diagram of the MPK6 crystal structure dimer that houses two copies of MPK6 in the asymmetric unit of 6DTL. Shown in yellow are the exposed Asp residues of the CD domain of MPK6 (D353, D356). **c,** Sequence alignment of the CD domain of AtMPK3/6 with animal MAPK orthologs (ERK and p38 family proteins) created using JALVIEW. The exposed residues of the CD domain are highlighted using red boxes. The CD domain exposed residues of MAPK are highly conserved in both plant and animal systems.

**Fig. S4:**
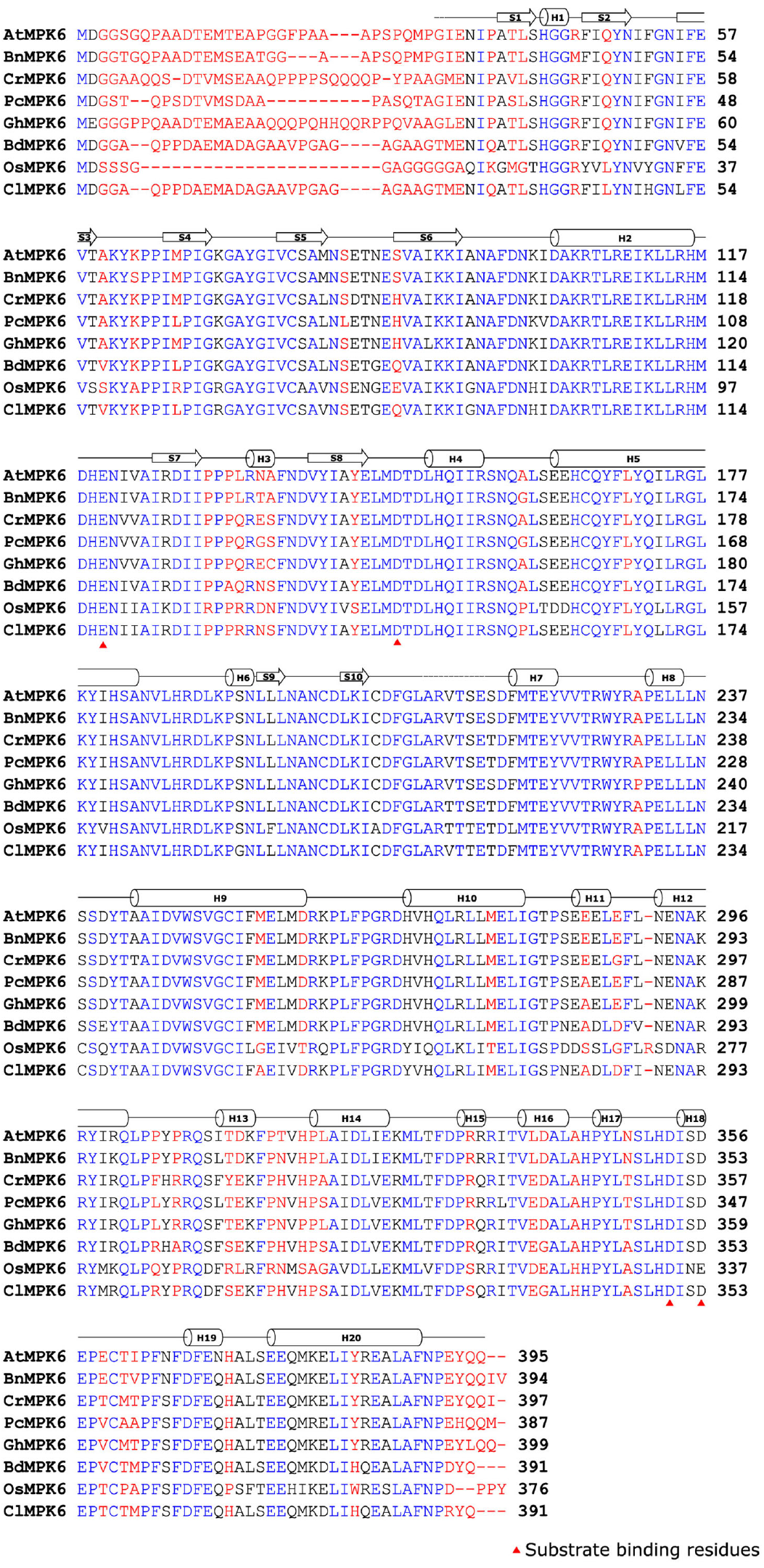
Secondary structure sequence alignment of AtMPK6. (Related to Figs. 4 and S3) Sequence alignment of the AtMPK6 along with its plant MPK6 orthologs. Secondary structure elements are depicted above the alignment (cylinder for α-helix and arrow for β-sheet). The exposed substrate binding resides on MPK6 (D148, E120, D353, D356) that are making contact with the SCRM KiDoK motif are depicted by red arrowheads.

**Fig. S5:**
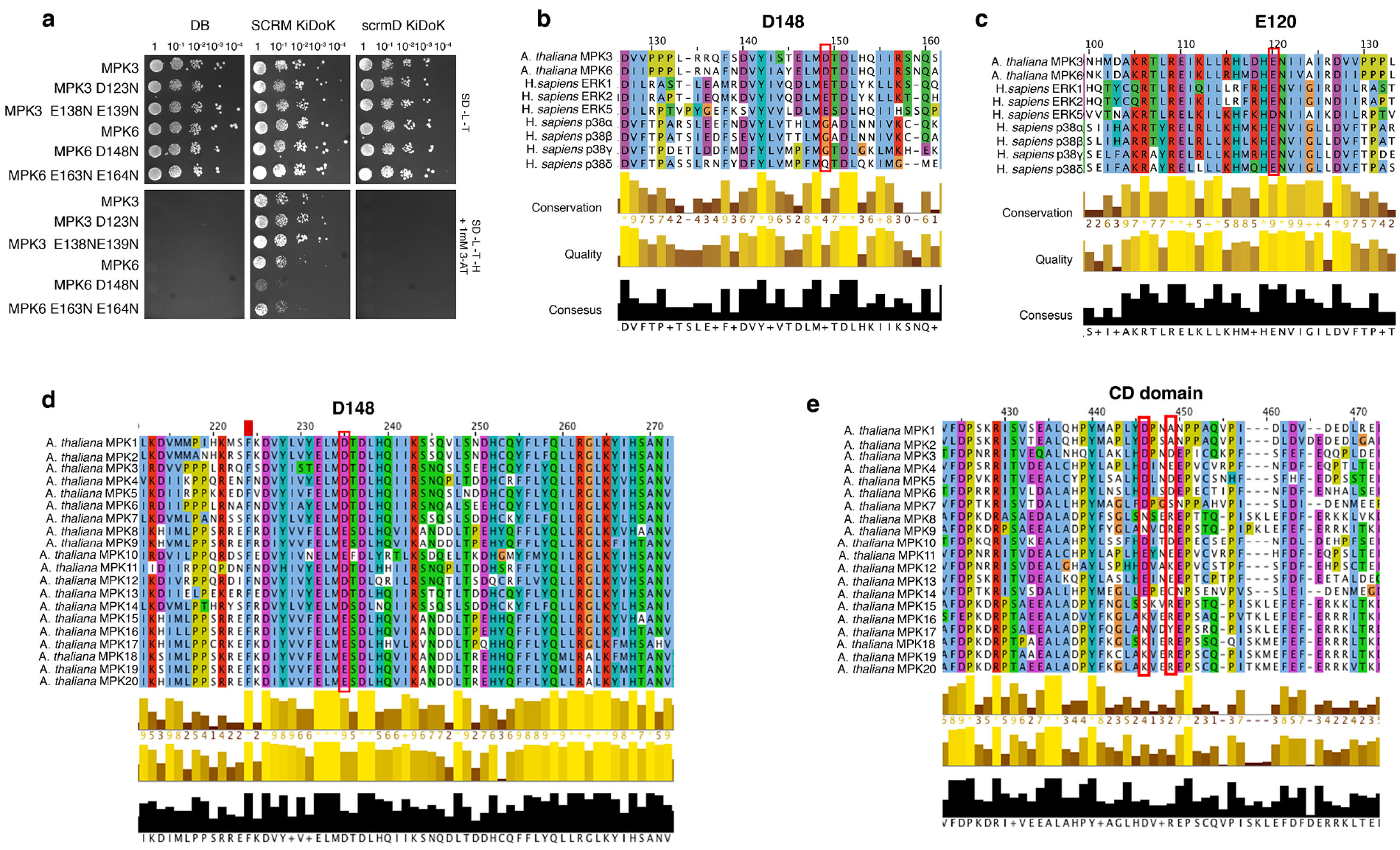
Interaction analyses and Sequence alignment of substrate binding residues within AtMPK6 (Related to Fig. 4) **a,** Y2H assays of SCRM-KiDoK and scrm-D KiDoK motifs with potential exposed substrate binding residues on AtMPK6 (identified using ab-initio modeling) mutated to Asn. Bait constructs containing the wild-type KiDoK motif (SCRM_KiDoK) and scrm-D version of the KiDoK motif (scrm-D_KiDoK) were tested in pairwise combinations with prey constructs containing the activation domain fused to MPK3, MPK3 D123N, MPK3 E138N E139N, MPK6, MPK6 D148N, MPK6 E163N E164N. Only MPK6 D148N showed severe reduction in interaction with the SCRM KiDoK motif, making D148 of MPK6 a substrate-binding site for the SCRM KiDoK motif. **b, c,** Sequence conservation analysis of the substrate binding residues, D148 and E120 of AtMPK6 against animal MAPK orthologs (ERK and p38 family proteins) created using JALVIEW. While E120 of AtMPK6 is conserved amongst both plant and animal MAPKs, D148 of AtMPK6 is far less conserved amongst animal MAPKs. **d, e,** Sequence conservation analysis of the substrate binding residues, D148 and D353, D356 of the CD domain of AtMPK6 amongst AtMAPK homologs (AtMPK1-20) created using JALVIEW. While D148 is highly conserved amongst all AtMAPK homologs, D353 and D356 of the CD domain of AtMPK6 are far less conserved amongst the other members of the AtMAPK family.

**Fig. S6:**
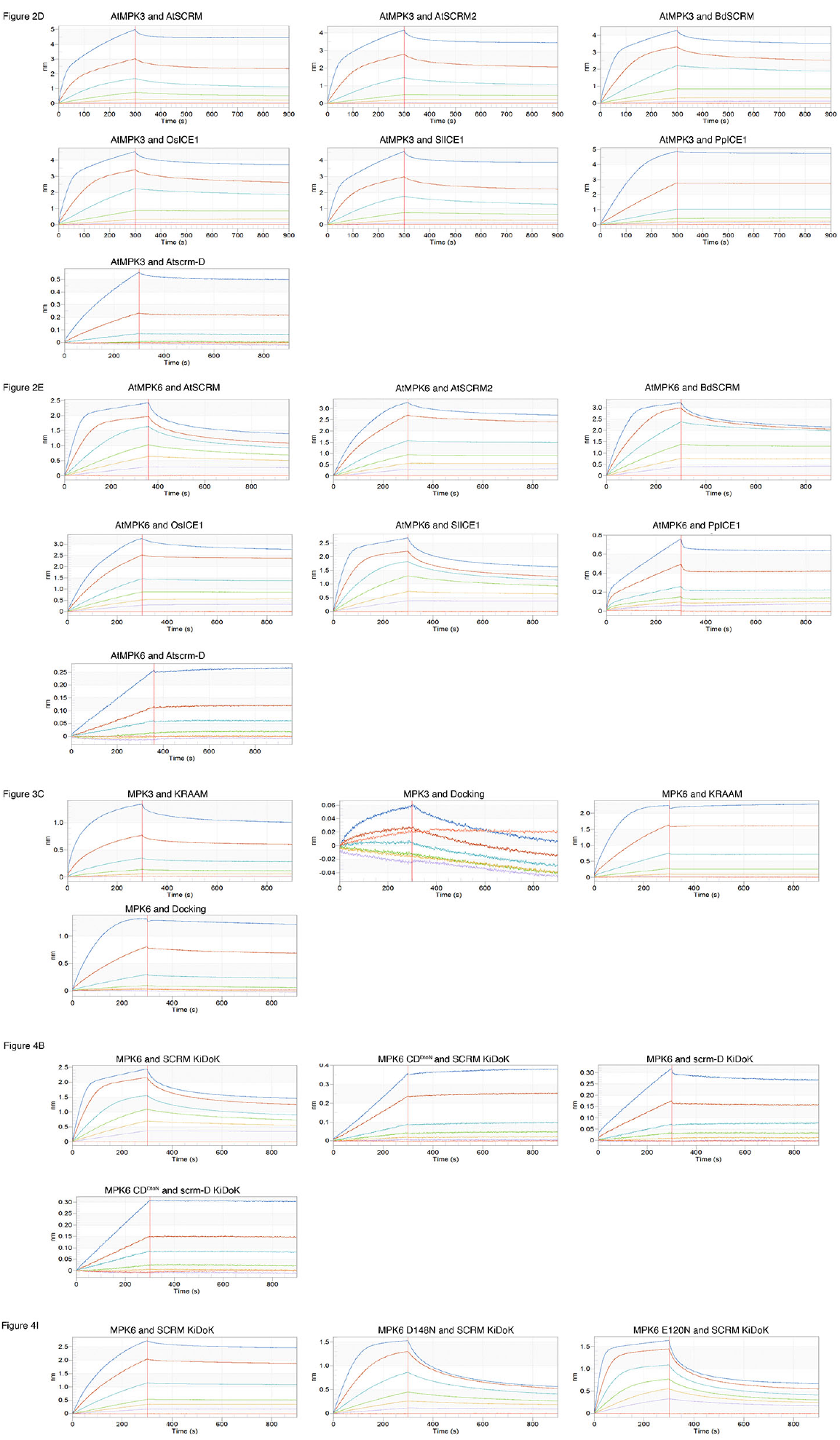
Raw sensogram traces of in vitro substate binding kinetics using the ForteBio Octet Red System (Related to Figs. 2, 3 and 4) Shown are representative raw sensogram traces of one (of three replicates) used in the *in vitro* binding kinetics experiments shown in Figs. 2,3 and 4. The 6 different concentrations used in every experiment are represented by blue, brown, cyan, green, yellow, purple and orange traces. Association with the peptide and protein was performed for the first 300 seconds followed by 600s of dissociation. The K_D_ values were calculated by fitting the association and dissociation curves with the 1:1 homogeneous ligand model.

